# Orderly specification and precise laminar deployment of cortical glutamatergic projection neuron types through intermediate progenitors

**DOI:** 10.1101/2024.03.01.582863

**Authors:** Dhananjay Huilgol, Jesse M Levine, William Galbavy, Bor-Shuen Wang, Z. Josh Huang

**Affiliations:** Department of Neurobiology, Duke University Medical Center; Durham, NC 27710, USA; Department of Biomedical Engineering, Duke University; Durham, NC 27708, USA; Cold Spring Harbor Laboratory; Cold Spring Harbor, NY 11724, USA; Program in Neuroscience and Medical Scientist Training Program, Stony Brook University; Stony Brook, NY 11794, USA; Program in Neuroscience, Department of Neurobiology and Behavior, Stony Brook University; Stony Brook, NY 11794, USA

**Keywords:** indirect neurogenesis, intermediate progenitors, neocortex, fate mapping, projection neurons (PNs)

## Abstract

The cerebral cortex comprises diverse types of glutamatergic projection neurons (PNs) generated from radial glial progenitors (RGs) through either direct neurogenesis or indirect neurogenesis (iNG) via intermediate progenitors (IPs). A foundational concept in corticogenesis is the “inside-out” model whereby successive generations of PNs sequentially migrate to deep then progressively more superficial layers, but its biological significance remains unclear; and the role of iNG in this process is unknown. Using genetic strategies linking PN birth-dating to projection mapping in mice, we found that the laminar deployment of IP-derived PNs substantially deviate from an inside-out rule: PNs destined to non-consecutive layers are generated at the same time, and different PN types of the same layer are generated at non-contiguous times. The overarching scheme of iNG is the sequential specification and precise laminar deployment of projection-defined PN types, which may contribute to the orderly assembly of cortical output channels and processing streams.

**HIGHLIGHTS:**

- Each IP is fate-restricted to generate a pair of near-identical PNs
- Corticogenesis involves the orderly generation of fate-restricted IP temporal cohorts
- IP temporal cohorts sequentially as well as concurrently specify multiple PN types
- The deployment of PN types to specific layers does not follow an inside-out order

## INTRODUCTION

The cerebral cortex contains a vast network of nerve cells underlying a wide range of high-level brain functions from sensory processing to cognition and motor control^1,2^. Across the horizonal axis of cortical surface, nerve cells are organized into dozens of cortical areas that form subnetworks of information processing^3,4^. Across the vertical axis of cortical thickness within each area, nerve cells are arranged in multiple cell layers and form intricate local circuitry^5^. Among the two major cortical neuron classes, whereas GABAergic interneurons extend local axons and regulate the balance and spatiotemporal patterns of cortical activity, glutamatergic projection neurons (PNs) elaborate local as well as long-range axons, linking local circuitry to global brain networks^5–7^. PNs constitute up to ~80% of cortical neurons and comprise diverse types characterized by their laminar location, morphology, projection patterns, and gene expression profiles^5,6,8–10^. The developmental mechanisms that specify these diverse PN types and organize them into characteristic cortical layers towards assembling the myriad cortical processing streams and output channels are not fully understood.

All PNs are generated from radial glia progenitors (RGs) lining the embryonic cerebral ventricle wall through two fundamental forms of neurogenesis^11,12^. In direct neurogenesis (dNG), a RG undergoes asymmetric division in the ventricular zone (VZ) to self-renew as well as generate one neuronal progeny^13–15^. In indirect neurogenesis (iNG), RG asymmetric division produces an intermediate progenitor (IP), which moves to the subventricular zone (SVZ) and undergoes symmetric division to generate two neurons^16–19^. Postmitotic PNs then undergo radial migration into the cortex, guided by radial fibers of RGs attached to the pia surface^20^. Classic cell birth dating experiments using tritiated thymidine in rodents^21^ and primates^22^ have revealed a systematic relationship between embryonic cell origin and their laminar location in the cortex. These seminal studies established a foundational concept of the inside-out order of corticogenesis, whereby successive generations of neurons migrate past earlier-generated cells to settle in progressively more superficial layers, forming a six-layered cortex in which a neuron’s laminar position is determined by its birth date^10,17,23,24^. However, these pioneering studies and their interpretation are limited by the coarse temporal and cellular resolution. First, subsequent higher temporal resolution studies using double S-phase labeling revealed that simultaneously born cells occupy more than one layer, and there is substantial overlap in the distributions of cells arising with successive cell cycles^25^. Second, the initial studies^21,22^ were carried out long before the discovery of the subpallium origin and tangential migration of GABAergic interneurons, thus did not distinguish the lamination of PNs from GABA interneurons^26^, which substantially complicates the interpretation of cell labeling and deployment pattern. Third, the findings were made long before the discovery of RGs and IPs as cortical neural progenitors^11,12,17,24^, and thus did not distinguish role of these progenitor types in PN production and laminar deployment. Therefore, to date, direct evidence for a stringent inside-out rule, whereby each successive generation of PNs migrate past their predecessor to generate progressively more superficial layers (as widely cited in most reviews^10,23,24^ and textbooks), has yet to be experimentally shown. Further, the biological significance of the inside-out order of cell settling in cortical development remains unclear, and the role of iNG in this process is unknown.

Between the two modes of neurogenesis, whereas dNG is ubiquitous along the neural tube that generate the entire central nervous system, iNG is restricted to the telencephalon that gives rise to the forebrain, especially the cerebral cortex^16,18,19,27–32^. Across evolution, while dNG originated before the dawn of vertebrates and has been conserved ever since, iNG emerged in amniotes and has expanded tremendously in mammals, driving the innovation of a six-layered neocortex^17,18,33–36^. Within the neocortex, whereas dNGs generate all major PN classes, iNG amplifies and diversifies PNs within each class^30^. Indeed, iNG generates the vast majority of PNs in neocortex (~80% in mice and even more in primates)^30,34,37,38^ that undergo the inside-out pattern of cortical migration. However, whether and how IPs and iNG contribute to the specification and laminar deployment of diverse PN types are unknown.

Here, combining a set of novel genetic strategies centered around TBR2, a transcription factor specifically expressed in IPs^29^, we fate map individual IPs and IP temporal cohorts with the resolution of individual cell division cycles. Furthermore, by integrating cell birth-dating with projection target mapping, we resolve the IP-derived PNs in terms of their morphology and projection targets much beyond laminar location. We found that individual IPs are fate-restricted to generate a pair of morphologically identical “twin” PNs. Surprisingly, the laminar deployment of successive IP-derived PNs substantially deviate from a simple inside-out rule: PN types destined to 2 or 3 non-consecutive layers are generated at the same time, and different PN types of the same layer are generated at non-contiguous times. Our results demonstrate that iNG involves the orderly specification of multiple temporal cohorts of fate-restricted IPs, which sequentially as well as concurrently generate numerous subsets of PN progeny, each deployed to a specific layer. These findings therefore suggest that, instead of a simple inside-out order of cell migration^17,21–24^, the overarching scheme of corticogenesis is the sequential specification and precise laminar deployment of projection-defined PN types, which may contribute to the orderly assembly of cortical output channels and processing streams.

## RESULTS

### Individual IPs are fate-restricted and generate pairs of near-identical PNs in the neocortex

Classic cell birth dating methods (e.g. tritiated thymidine, BrdU) are limited by their lack of specificity to progenitor types (e.g. pallium GLU vs subpallium GABA progenitors, RGs vs IPs) and cannot resolve the phenotypes of birth-dated cells beyond their location. As the majority of cortical intermediate progenitors (IPs) in mice undergo a single round of symmetric neurogenic division, and transit-amplifying IPs are estimated to be only ~10%^16,19,27,39^, genetic targeting of IPs is a specific and precise fate mapping approach to iNG-derived PNs. We have generated a *Tbr2-2A-CreER* driver in which a *2A-CreER* cassette is inserted at the end of the *Tbr2* coding sequence, thereby preserving the gene function^40^. Extensive characterization of this driver line demonstrates that it achieves specific targeting of IPs. Indeed, short (12 hr) pulse-chase experiments by tamoxifen (TM) induction in *Tbr2-CreER;Ai14* mice at embryonic days E13.5, E15.5 and E16.5 labeled only multipolar cells in the subventricular zone (SVZ) but not RGs in the ventricular zone (VZ) (Figure 1a-f). Most of these SVZ multipolar cells were immune-positive for TBR2 (62-85%) throughout corticogenesis (Figure 1e) and for the cycling progenitor marker, Ki67 (~65%) at early and late stages of neurogenesis (Figure 1f). Other RFP^+^ cells labeled at these stages were likely postmitotic cells, such as Cajal-Retzius cells and olfactory projection neurons migrating to the accessory olfactory bulb (Figure 1f)^32,41^. Longer (48 or 72 hr) pulse-chase at E11.5 or at E14.5 did not label any persisting progenitors in the germinal zone (Figure S1) indicative of potential proliferative basal radial glial cells (bRGs)^34,42^, consistent with IPs having a cell cycle length of around 24 hours^43^. Therefore, the *Tbr2-2A-CreER* driver specifically targets IPs.

**Figure 1:**
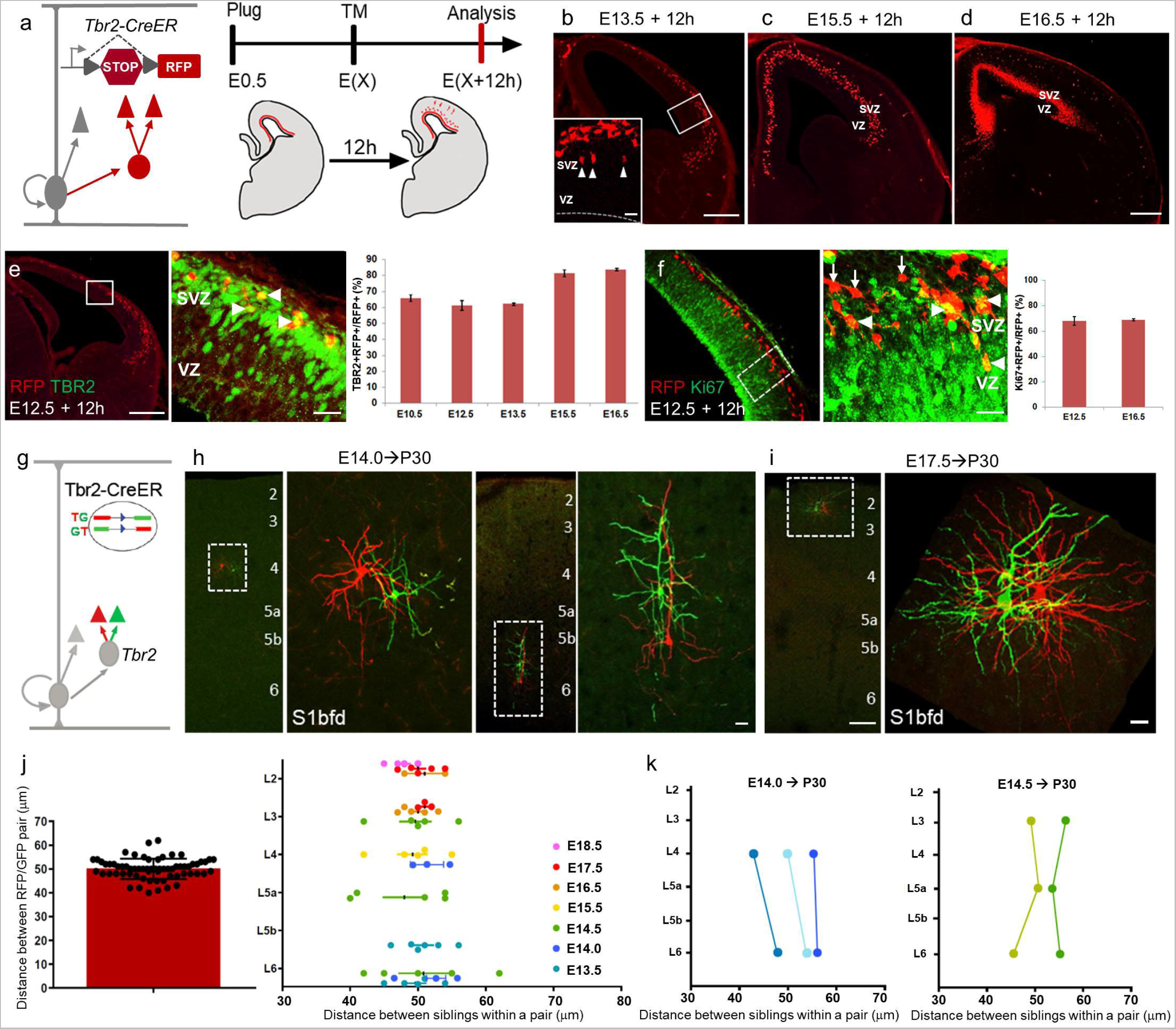
Individual IPs are fate-restricted to generate a pair of near-identical twin PNs. (a) Genetic strategy using Tbr2-2A-CreER and a Cre-dependent reporter Ai14 to label IPs and iNG-derived PNs (red). Experimental scheme (right) for short pulse-chase fate mapping where tamoxifen (TM) induction at different embryonic times is analyzed 12 hours post-induction. (b) Coronal hemisection of a Tbr2-CreER;Ai14 embryonic brain at E13.5 following a 12hr TM induction reveals a lateral-high to media-low gradient of RFP+ cells. High magnification inset shows RFP-labeled cells away from the VZ with round or multipolar morphology indicating IPs (arrowheads). (c,d) Tbr2-Ai14 coronal hemisections at E15.5 (c) and E16.5 (d) show labeling of cells in the subventricular zone (SVZ) similar to E13.5 but with no apparent gradient. (e) Coronal hemisections similar to (b-d) showing TBR2 immunolabeling (green) on Tbr2-CreER;Ai14 embryonic brains at E12.5 following a 12hr TM induction. Colocalization of RFP and TBR2 (arrowheads) are quantified at different embryonic times (right). (n=6 embryos, 2 litters at each age). (f) Coronal hemisection similar to (e) shows cycling progenitor marker Ki67 colocalized (arrowheads) and non-colocalized (arrows) with RFP; colocalization at E12.5 and E16.5 are quantified (right). (n=5 embryos, 2 litters at both ages). (g) Schematic for clonal analysis of IPs using Tbr2-2A-CreER;MADM mice (Mosaic analysis with Double Markers). Cells derived from the same cell division are differentially labeled by RFP and GFP. (h) E14 TM induction labeled morphologically near identical pairs of RFP+ and GFP+ stellate neurons in L4 (left) and PNs in L6 (right). (n=5 animals, 3 litters). (i) TM at E17.5 labeled near identical pair of L2 PNs. (n = 6 animals, 3 litters) (j) Distance between twin cells generated from IPs at different ages averaged across all 80 pairs (left) or separated by their age of induction and laminar positions (right). (n = 5-6 animals, 2-3 litters each age). (k) Concurrent labeling of twin PNs across non-consecutive layers in the same cortical area in mice induced at E14.0 and E14.5. PN twin pairs in the same cortical area from the same animal are connected by a line. 2 of 5 animals (TM E14.5) and 3 of 6 animals are represented (TM E14.0). Scale bars, low mag images = 100μm, high mag images = 20μm. Abbreviations: VZ, ventricular zone; SVZ, subventricular zone; TM, tamoxifen; S1_bfd_, primary somatosensory barrel field cortex. See also Figure S1, Supp video S1.

While the vast majority of IPs in mice undergo a single round of symmetric cell division to generate two PNs^16,19,27^, it remains unclear to what extent the two PN progeny are phenotypically similar or different, i.e., whether they are of the same or different cell type. We combined *Tbr2-2A-CreER* driver with the clonal analysis *MADM* reporter (mosaic analysis using double marker)^44,45^ to compare the two differentially labeled (RFP or GFP) PN progeny derived from individual IPs (Figure 1g). As the frequency of inter-chromosome recombination of *MADM* is very low, TM induction from E13.5 through E18.5 in 35 *Tbr2-2A-CreER;MADM* mice generated a total of 80 very sparsely labeled pair of PNs in the neocortex analyzed at ~P30. Strikingly, the two PN progeny within each pair were always located in close proximity within the same layer and showed highly similar dendritic morphology (Figure 1h-j). This result suggests that individual IPs are fate-restricted to generate a pair of “twin PNs” of the same laminar and morphological type. Notably, TM induction at E14.0 gave rise to pairs of spiny stellate cells in L4 as well as a pairs of L6 PNs, and E14.5 induction gave rise to pairs of L6, L5a, and L3 PNs all within the same area (Figure 1j,k); these results suggest the simultaneous presence of two or more cohorts of fate-restricted IPs at this stage in the germinal zone. During late phase corticogenesis, E17.5 TM induction labeled twin PNs in L2 (Figure 1i). Of the total of 80 PN pairs examined, the distance between the two GFP and RFP labeled PNs within each pair was on an average ~50um (Figure 1j,k; Supp video S1). Together, these results suggest that individual IPs are fate-restricted to generate pairs of PNs of the same type, and the *Tbr2-2A-CreER* driver achieves fate mapping of iNG-derived PNs with the temporal resolution of a single cell division cycle.

### Orderly and precise laminar deployment of IP-derived PNs substantially deviates from a simple inside-out pattern

While highly informative, the exceedingly low frequency of the *MADM* reporter precluded an efficient fate mapping of IPs. We therefore used the *Tbr2-2A-CreER;Ai14* mice to carry out a comprehensive fate mapping of IPs throughout corticogenesis at each embryonic day from E11.5 to E18.5 (Figure S2). As iNG generates PNs for all pallium-derived cortical regions^30,31^, we first examined the global patterns of PN production across broadly defined cerebral cortex structures. TM induction from E11.5 through E18.5 revealed a lateral-to-medial gradient of iNG, reflecting the embryonic gradient of *Tbr2* expression (Figure 1b,e). The piriform cortex PNs were generated earliest from E11.5 through E15.5, followed by the basolateral amygdala and the insula at E12.5 and E13.5. Hippocampal PN production had a late onset at E14.5 in CA1 and CA3, followed by DG subfields at E15.5; CA1 and DG neurons continue to be born from E16.5 through E18.5 (Figure S2). Neocortical PNs were generated during the longest time window from E11.5 through E18.5, broadly following a lateral-to-medial areal gradient and deep-to-superficial layer order (Figure S2).

To further characterize the pattern of iNG-mediated PN production and their laminar deployment in the neocortex, we focused our analysis in the primary somatosensory cortex barrel field (S1_bfd_) across the comprehensive fate mapping time series (Figure 2a, b). TM induction at E12.5 labeled mostly L6 PNs at P30 and E13.5 induction labeled PNs in L6, L5b and L5a, consistent with an inside-out sequence of PN production during the early phase of neurogenesis. Surprisingly, TM induction at E14.0 produced a striking pattern, generating PNs predominantly in L4 and L6 but not in L5. This result corroborated the one observed at the level of individual IPs using the *MADM* reporter (Figure 1h,k), together suggesting that there appeared to be two populations of fate-restricted IPs at this stage giving rise to PNs destined to non-consecutive layers. Subsequently, in a span of 12 hours during this phase of peak neurogenesis, E14.5 TM induction generated an even more unexpected lamination pattern, where PNs in L6, L5a, and L3 but not L5b and L4 were labeled. This result again corroborated the *MADM* reporter data (Figure 1k), and suggests the presence of 3 fate-restricted IP populations simultaneously giving rise to PNs deployed to 3 non-consecutive layers. Further deviating from a simple inside-out order, TM induction at E15.5 labeled L4 PNs again, which were produced previously at E14.0; and E16.5 induction generated L3 PNs again, which were previously produced at E14.5. Therefore, both L4 and L3 PNs were produced from at least two non-contiguous developmental times or cell division cycles. During late phase neurogenesis, E17.5 induction generated L3 and L2 PNs, and E18.5 induction labeled the most superficial L2 PNs. However, E18.5 induction also labeled a set of L5a PNs, which constituted approximately 10% of the population generated at this time (Figure 2b). Therefore, IP-mediated PN production and deployment in the cortex substantially deviates from the inside-out model, including generating PNs in non-consecutive layers at the same time, generating PNs of the same layer at non-contiguous times, and ending PN production with an “outside-in” pattern.

**Figure 2:**
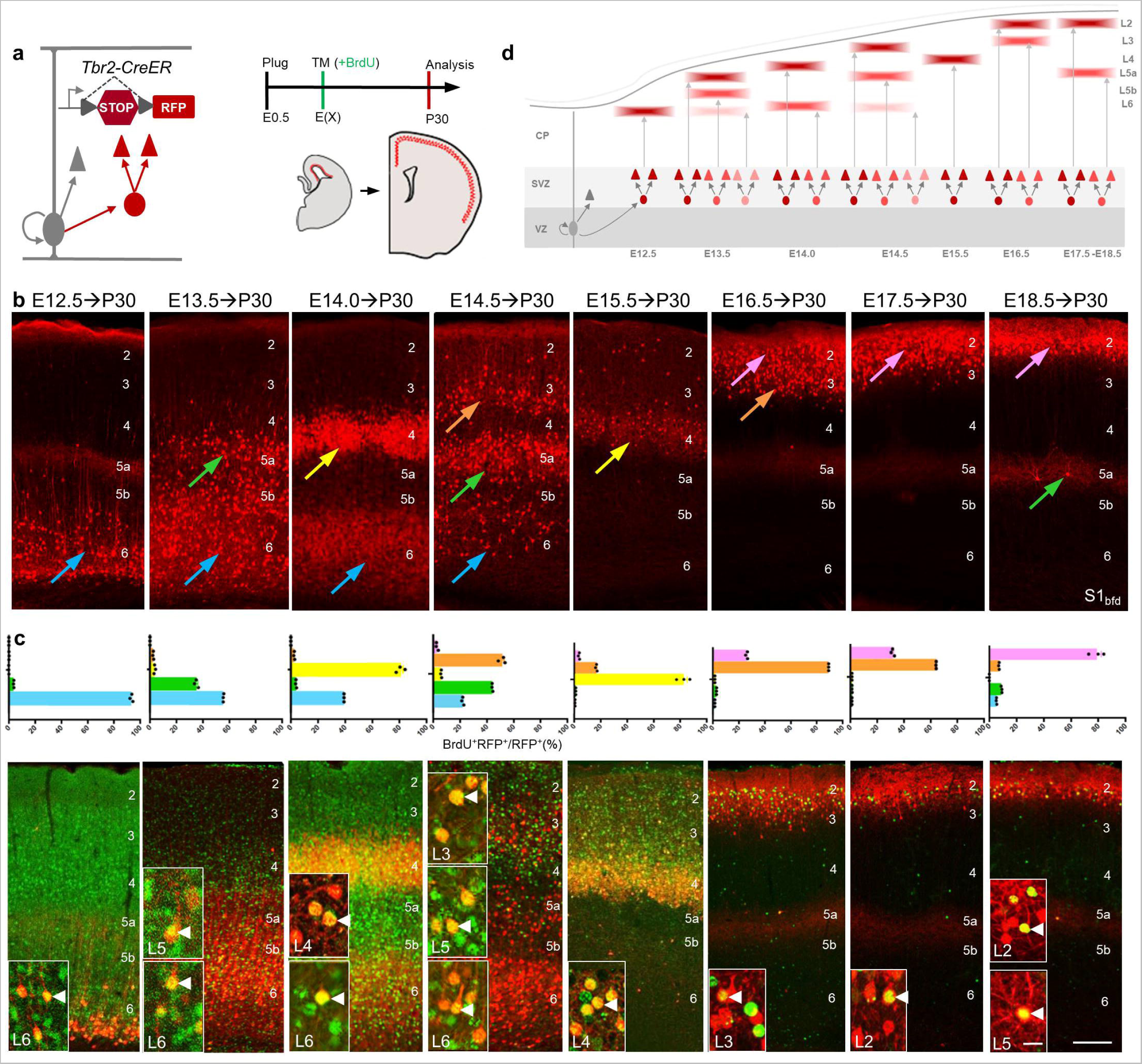
Orderly generation and precise laminar deployment of IP-derived PNs substantially deviates from a simple inside-out lamination pattern. (a) Experimental scheme of TM induction in Tbr2-2A-CreER;Ai14 embryos (left) or TM co-administered with BrdU (green) at different embryonic times followed by analysis at P30. (b) Lamination patterns of IP-derived PNs generated across embryonic times (as indicated) show an overall inside-out trend. Note that E14-born PNs occupy L6 (blue arrow) and L4 (yellow arrow), E14.5-born PNs occupy L6, L5a (green arrow), and L3 (orange arrow). L4 PNs are generated at separate times (E14 and E15.5). L3 PNs are generated at both E14.5 and E16.5 (n = 6-7, 3 litters). E18.5 induction labeled a small fraction of L5a (10%, green arrow) in addition to L2 (pink arrow) PNs (n=6, 2 litters). (c) Co-administration of BrdU with TM at each embryonic time point validates the correlation between PN birth time and lamination pattern shown in (b). Quantification of percentage BrdU+/RFP+ of total RFP+ cells (top row) showed labeling similar to Tbr2-CreER;Ai14 (b). Representative images (bottom row) show BrdU+ and RFP+ PNs at each time point. Insets show colocalization of RFP+BrdU+ PNs (arrowheads). (n = 5, 2 litters at each age). (d) Summary schematic showing that iNG involves the orderly specification of multiple temporal cohorts of fate-restricted IPs, which sequentially as well as concurrently generate numerous subsets of PN progeny, each deployed to a specific layer; as a result, the laminar deployment of successive IP-derived PN subpopulations substantially deviate from a simple inside-out rule. Scale bars, low mag images showing cortical layers = 100μm, high mag images = 20μm. Abbreviations: S1_bfd_, primary somatosensory barrel field cortex, BrdU, bromodeoxyuridine. See also Figures S2, S3.

To substantiate results obtained from the *Ai14* reporter, we employed the *RGBow* reporter^40,46^, which features lower frequency *Cre* recombination and allows sparser labeling of IPs and PNs (Figure S3). Low dose TM induction in *Tbr2-CreER;RGBow* mice at E14.0, E14.5 and E15.5 confirmed the production of PNs in non-consecutive layers (i.e., L4 and L6 at E14.0, and L6, L5a, L3 at E14.5) and the production of PNs of the same layer at non-consecutive days (i.e., L4 from E14.0 and E15.5, and L3 from E14.5 and E16.5; Figure S3).

To further validate the *Tbr2-2A-CreER;Ai14* based birth dating method, we co-administered bromodeoxyuridine (BrdU), a well-established cell birth-dating marker, along with TM induction to ensure that the observed laminar patterns accurately captured cell birth time (Figure 2c). Colocalization analysis of BrdU^+^ and RFP^+^ PNs from E12.5 through E18.5 confirmed the birth pattern of PNs (Figure 2c). Indeed, L4 and L6 PNs were co-labeled by TM and BrdU administration at E14.0, and L3, L5a and L6 PNs were co-labeled by TM and BrdU at E14.5 (Figure 2c). Notably, TM and BrdU administered at E18.5 resulted in co-labeling of not only L2 but also L5a PNs, confirming the late birth of deep layer PNs (Figure 2c).

To examine whether the pattern of multiple non-consecutive layers could be due to generation of PNs from latent or transit-amplifying IPs^16^, we incorporated another birth dating marker, EdU, 24-hours after TM induction to capture potential transit-amplifying IPs. Less than 4% and 2% RFP^+^ PNs were co-labeled with EdU at E14.0 and E14.5, respectively. This confirmed that the vast majority of IP-derived PNs were born less than 24-hours of TM induction (Figure S3), indicating that PNs residing in non-consecutive layers have the same birthdate. Furthermore, we used an independent genetic method to specifically target the neurogenic (i.e. not the transit-amplifying) set of Tbr2^+^ IPs by intersection with TIS21, an anti-proliferative protein that is specifically expressed in neurogenic progenitors^47^. We combined our previously generated inducible *Tis21-CreER* and *Tbr2-FlpER* drivers^40,48^ with a *Cre-* and *Flp-*intersectional reporter, *Ai65*^49^ to restrict birth dating of PNs derived from neurogenic IPs. TM induction at E14.0 again generated PNs in L4 and L6 (Figure S3), corroborating the results from *Tbr2-CreER;Ai14* fate mapping (Figure 2b) as well as from the BrdU experiments (Figure 2c). Both these results were also consistent with data from *Tbr2-CreER;MADM* mice induced at E14.0, which labeled ‘twin’ PNs in L4 and L6 (Figure 1h,k) Together, these results indicate that the simultaneous generation of PNs destined to non-consecutive layers results from concurrent neurogenesis from two or more cohorts of fate-restricted IPs.

Taken altogether, our results suggest that cortical iNG is characterized by the sequential specification of multiple temporal cohorts of fate-restricted IPs, each generating a subpopulation of PNs that is precisely deployed to a specific cortical layer. The overall lamination patterns of these orderly as well as concurrently generated PN laminar subsets substantially deviate from a simple inside-out order (Figure 2d).

### Early IP-derived PNs include deep layer ET neurons

Beyond laminar location, a defining feature of PN type identity is their axon projection targets, but most previous fate mapping methods do not resolve the anatomical properties of birth-dated PNs. We therefore designed a novel genetic intersection strategy that integrates birth-dating and projection mapping of IP-derived PNs. We combined *Tbr2-2A-CreER* driver with an *intersection-subtraction (IS)* reporter, which expresses RFP upon *Cre* activation and GFP upon *Cre*(AND)*Flp* activation^46^. In *Tbr2-2A-CreER;IS* mice (Figure 3a), embryonic TM induction of *CreER* marks the birth time of IP-derived PNs with RFP, and subsequent postnatal injection of a retrograde *AAV-Flp*^50^ to a brain region activates GFP expression in the subset of birth-dated PNs that project to the injection site. For comparison, co-injection of a generic retrograde FluoroGold tracer with *retroAAV-Flp* labels all PNs projecting to the target. We applied this strategy at each embryonic day across corticogenesis.

**Figure 3:**
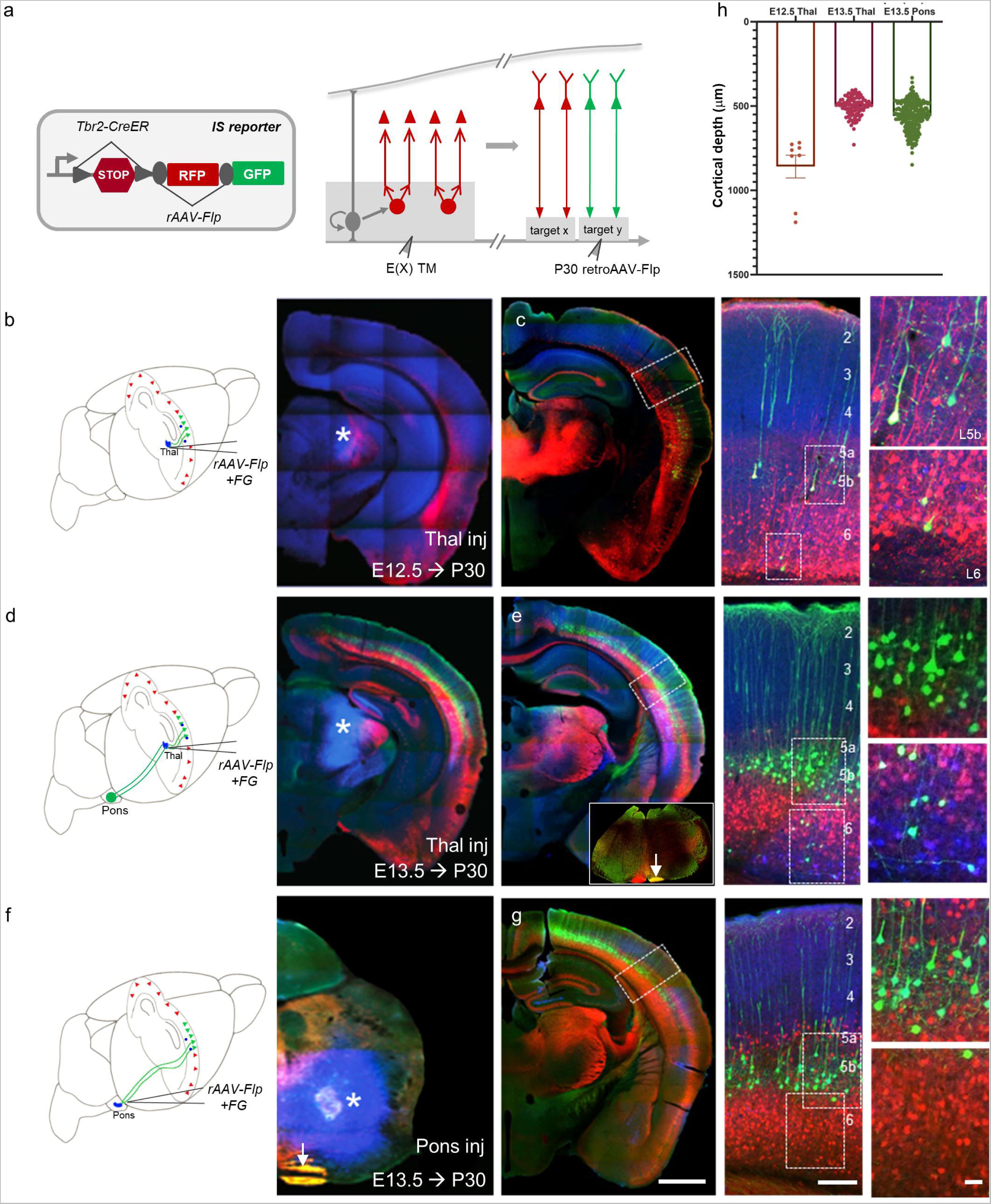
Early iNG-derived PNs include L6 CTs and L5b PTs. (a) A genetic fate mapping strategy of PNs that integrates their IP origin, birth date, and projection targets. Using an intersection-subtraction reporter (IS) bred with Tbr2-CreER driver (left), embryonic TM induction marks the birth time of IP-derived PNs with RFP expression; postnatal injection of rAAV-Flp then converts RFP to GFP expression in PNs that project to the injection target (right). Co-injection with fluorogold (blue) provides a generic label to all PNs projecting to the injection target. (b) Schematic (left) and coronal image (right) showing AAV injection into the VPM/Po nuclei of the thalamus (asterisk) in P30 Tbr2-CreER;IS mouse TM-induced at E12.5. (c) Among E12.5 born PNs (RFP^+^) in S1bfd, a subset in L5b and L6 projected to the thalamus (GFP^+^) (higher magnification views of boxed regions shown in right panels). (d,e) Thalamic injection scheme (d) in a mouse TM induced at E13.5 (asterisk, right panel). Among RFP^+^ E13.5 born L5/6 PNs, a large set of L5b and sparse set of L6 PNs projected to thalamus (GFP^+^, e); higher magnification views of boxed regions shown in right panels). Inset in coronal section (e) shows GFP^+^ axon collaterals in the pons (arrow). (f,g) Pons injection (f) in a P30 mouse TM induced at E13.5 (asterisk, right). Among RFP^+^ E13.5 born L5/6 PNs, L5b PNs projected to pons (high magnification views shown in right panels) (g). Note GFP^+^ axon tract in the pons (arrow in f). (h) Quantification of cortical depth of GFP^+^ PNs with thalamus rAAV-Flp injection in mice TM-induced at E12.5 (brown) and E13.5 (red), and with pons rAAV-Flp injection in mice TM-induced at E13.5 (green). The majority of GFP-labeled PNs were in L5b. Asterisk, site of injection (b,d,f). Scale bars = 1mm for brain hemisections, low mag images showing cortical layers = 100μm, high mag images = 20μm. Abbreviations: IS, intersection-subtraction; rAAV, retrograde adeno-associated virus; FG, fluorogold; Thal, thalamus. See also Figure S4.

Extratelencephalic (ET) neurons, including the corticothalamic (CT) and pyramidal tract (PT) neurons are born during early corticogenesis^10^. We thus used *rAAV-Flp* injection to the thalamus and pons in *Tbr2-2A-CreER;IS* mice induced at E12.5 and E13.5 to examine the production of these PNs. *rAAV-Flp* injection to the thalamus or pons mainly in E13.5-induced and sparsely inE12.5-induced mice labeled a large set of PNs in L5b, which extended prominent and characteristic apical dendrites to the pia and projected axons along the pyramidal tract to pons and beyond (brainstem). This result indicates that IP-derived PT neurons are generated at ~E13.5 (Figure 3b-h).

In contrast, *rAAV-Flp* injection to the thalamus in E12.5-induced mice labeled only a sparse set of L6 neurons, even though the co-injected FluoroGold labeled a broad set of L6 cells (Figure 3b,c,h). To ensure that the sparse labeling by *rAAV-Flp* was not due to technical factors such as the limited coverage of target area, we performed a broad thalamus injection with another generic retrograde label CTB^647^ in the *Tbr2-Flp;FSF-tdTomato* mice (a *Flp*-dependent reporter) in which all iNG-derived PNs were RFP labeled irrespective of their birth time^30,40^. Quantification showed that 18.8% CTB^+^ cells colocalized with RFP^+^ cells in L6 across different cortical areas (Figure S4), consistent with our previous finding that a minor but significant proportion of L6 CT PNs were generated from IPs^30^. Together, these results suggest that early IP-derived PNs include a small set of L6 CT and major set of L5b PT neurons.

### IP-derived corticostriatal and callosal PNs laminate in a largely inside-out order

The intratelencephalic (IT) PNs include diverse cell types located across all layers and are generated throughout corticogenesis. Among these, two major subpopulations are the corticostriatal (CSPN) and callosal (CPN) PNs, which show extensive overlap^10,51,52^. To birthdate IP-derived CSPNs, we injected *rAAV-Flp* to the dorsolateral striatum (DL-Str), the striatal target region for PNs in S1_bfd_, in *Tbr2-2A-CreER;IS* mice TM-induced at embryonic days across corticogenesis. In E13.5-induced mice, DL-Str injection labeled cells in L6, L5b, and L5a (Figure 4a,b,f); a subset of labeled L5b cells may include PT neurons which extend collaterals to the striatum (Figure 3f,g)^53–55^. DL-Str injection in E14.5- and E16.5-induced mice labeled a large set of cells in L5a and L3, respectively (Figure 4c,d,f). Finally, DL-Str injection in E18.5-induced mice labeled a small set of cells in L2 (Figure 4e,f). Together, these results suggest that IP-derived CSPNs are generated throughout mid-to late corticogenesis in a largely inside-out order.

**Figure 4:**
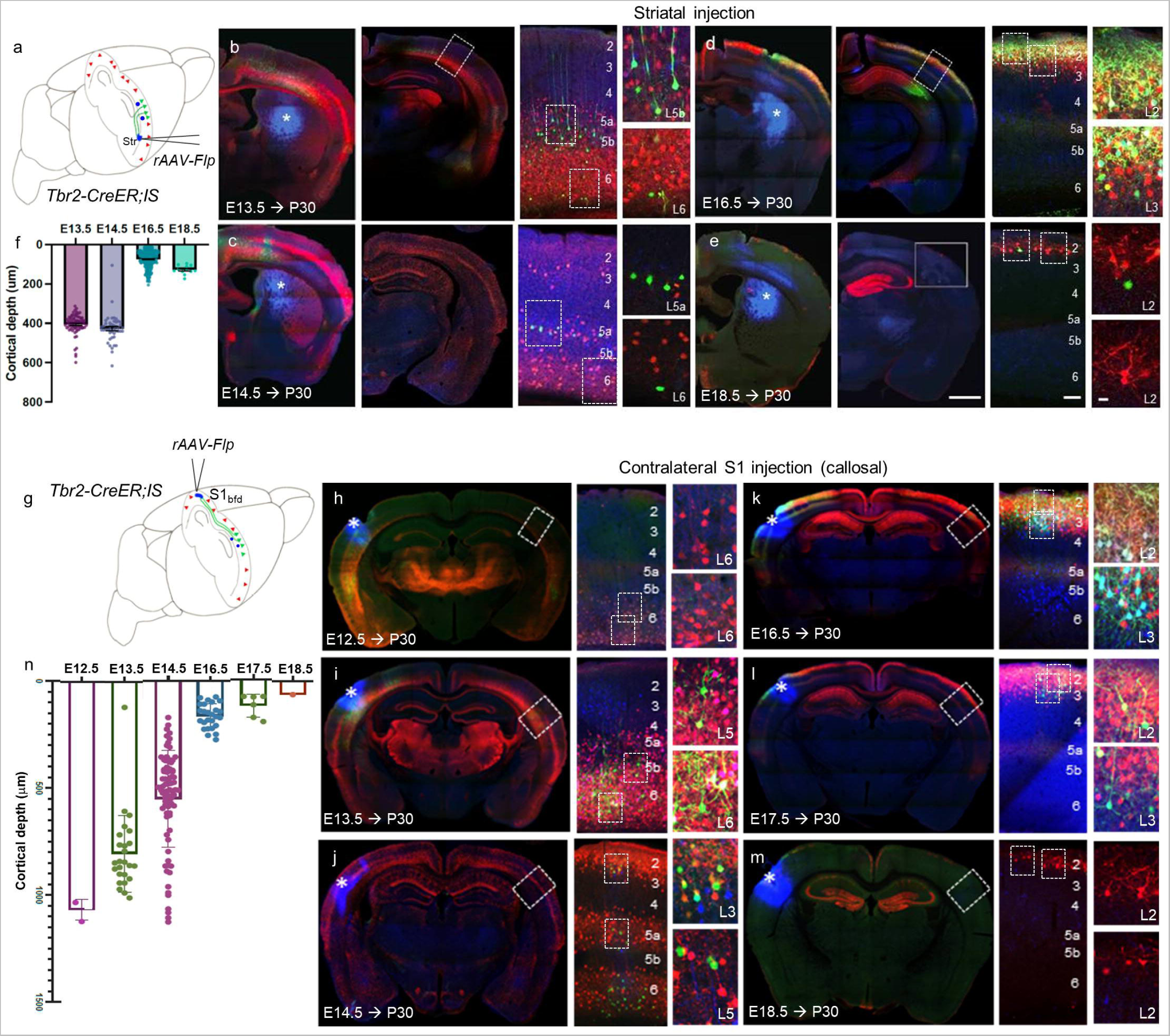
IP-derived corticostriatal and callosal ITs are specified sequentially from deep to superficial layers. (a) rAAV injection strategy for fate mapping striatum-projecting PNs in Tbr2-CreER;IS mice. (b) rAAV-flp injection (asterisk, left) revealed that among E13.5 born PNs (RFP^+^) in S1bfd, a subset in L5a, L5b and L6 projected to the striatum (GFP^+^) (higher magnification views of boxed regions, right panels). (c) Among E14.5 born PNs in L3, L5a and L6 (RFP^+^), L5a and a small fraction of L6 PNs (GFP^+^) project to the striatum (d) A large fraction of L2 and L3 PNs TM-induced at E16.5 (RFP^+^) are striatal-projecting (GFP^+^). (e) Some superficial L2 PNs TM-induced at E18.5 (RFP^+^) project to the striatum (GFP^+^). (f) CSPNs are broadly generated in a deep-to-superficial order. Each dot represents a GFP^+^ PN. Y axis = cortical depth (μm). (g) Same strategy as in (a) for fate mapping PNs projecting to contralateral cortex. (h) E12.5 born PNs (RFP^+^) in S1bfd did not project to contralateral S1bfd (absence of GFP^+^ PNs) (i) rAAV-flp injection in S1bfd of mice TM-induced at E13.5 labeled a set of L5b/6 PNs (GFP^+^), indicting their projection to contralateral S1bfd among PNs born at E13.5 (RFP^+^) (j) Among E14.5 IP-derived PNs (RFP^+^) in S1bfd, subsets in L3, L5a and L6 (GFP^+^) project to contralateral S1bfd. (k,l) Among IP-derived PNs TM-induced at E16.5 (k) and E17.5 (RFP+), a large subset in L3 (GFP^+^) projected to contralateral S1. (m) E18.5 IP-derived PNs (RFP^+^) did not project to the contralateral S1bfd (i.e. no GFP labeling). (n) GFP+ CPNs (dots) in S1bfd born at different embryonic times show an overall inside-out pattern of laminar distribution. X axis: age of TM induction, Y axis: depth from pia (μm). High magnification of boxed areas are shown in right panels of b-e, h-m. Asterisk, site of injection (b-e, h-m). Scale bars = 1mm for brain hemisections, low mag images showing cortical layers = 100μm, high mag images = 20μm. Abbreviations: IS, intersection-subtraction; rAAV, retrograde adeno-associated virus; S1_bfd_, primary somatosensory barrel field cortex; Str, striatum.

To birthdate IP-derived CPNs, we injected *rAAV-Flp* to the contralateral S1_bfd_ (cS1) in *Tbr2-2A-CreER;IS* mice TM-induced from E12.5 to E18.5, and examined cell labeling patterns in the ipsilateral S1_bfd_ (iS1) (Figure 4g-n). *rAAV-Flp* injection in E12.5- and E18.5-induced mice showed only sparse or no GFP labeling in iS1, suggesting that CPNs are not generated from IPs at the beginning and end of corticogenesis (Figure 4g,h,m,n). *rAAV-Flp* injection in mice induced between E13.5 and E17.5 labeled a large set of CPNs in iS1 located progressively from deep layers (L6 and L5b in E13.5-induced, L6, L5a, and L3 in E14.5-induced) to more superficial layer (L3 and L2 in E16.5-induced, L2 in E17.5-induced). Notably, *rAAV-Flp* injection in E14.5-induced mice labeled CPNs across non-consecutive layers (L6, L5a, and L3), suggesting the presence of at least 3 fate-restricted cohorts of IPs simultaneously generating 3 subtypes of CPNs at this peak time of neurogenesis (Figure 4j,n).

### Intra-hemispheric PNs are generated early for deep layers and late for superficial layers

Another major subpopulation of IT neurons is the intrahemispheric-projecting PNs (IPNs) that restrict their axons to the same hemisphere^10,51,52^. To birthdate IP-derived IPNs, we injected *rAAV-Flp* to M2 in *Tbr2-2A-CreER;IS* mice TM-induced at embryonic days across corticogenesis and examined cell labeling patterns in S1_bfd_ of the same hemisphere (iS1) (Figure 5). Overall, the cell labeling patterns in S1_bfd_ in mice induced from E12.5 to E18.5 progressed from deep to superficial layers with an order similar to that of CPNs (Figure 4n), but with at least two notable differences. First, whereas CPNs were hardly generated at E12.5 and E18.5, IPNs were generated at the earliest as well as the latest times of corticogenesis. Second, while *rAAV-Flp* injection in cS1 in E16.5-induced labeled a significant proportion of CPNs in L3 of iS1 (Figure 4k,n), iM2 injection at the same age labeled neurons more in L2 than L3 of iS1, suggesting a distinction between L3 CPN and L2 IPNs (Figure 5e,h). Again, IPNs in 2 or more layers were labeled in E13.5- and E14.5-induced mice (Figure 5c,d,h), suggesting the presence of multiple fate-restricted IP cohorts at these stages. As M2 *rAAV-Flp* injection would label all PNs with an axon branch to M2, possibly including CSPN and CPN, the results from E14.0 and E14.5 induced mice do not specifically fate map IPNs^51,52^.

**Figure 5:**
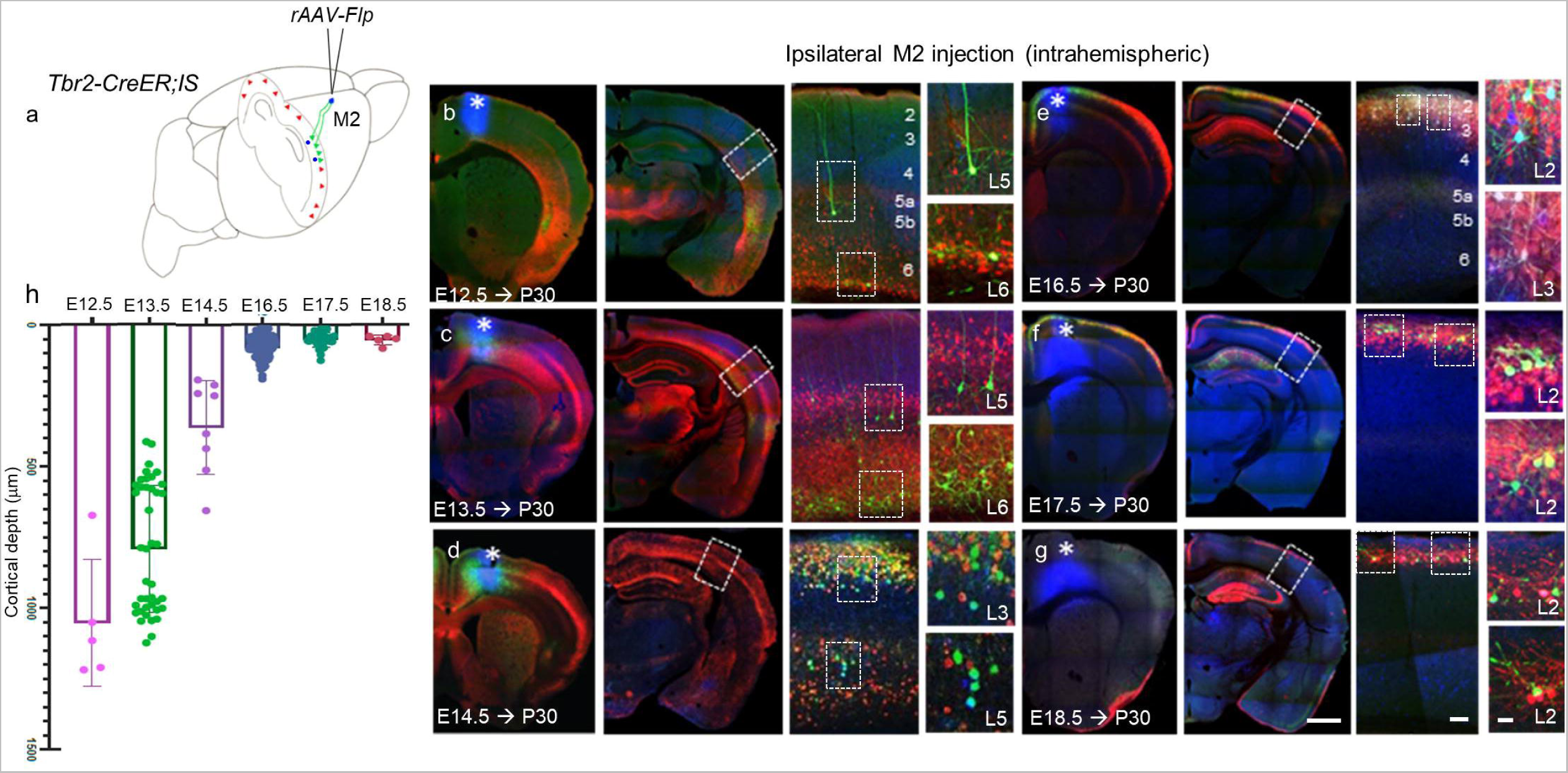
IP-derived intrahemispheric ITs are generated at the earliest and last phases of indirect neurogenesis. (a) rAAV injection strategy for fate mapping IPNs in Tbr2-CreER;IS mice. (b,c) Among IP-derived PNs in S1bfd born at E12.5 (b) and E13.5 (c) (RFP^+^), a subset in L5/6 expressed GFP following rAAV-flp injection to ipsilateral M2 (iM2). (d) Among IP-derived PNs in S1 TM-induced at E14.5 (RFP^+^), subsets in L3 and L5a GFP^+^ projected to iM2. (e,f,g) Among IP-derived PNs (RFP^+^) in S1 bfd TM-induced at E16.5 (e), E17.5 (f) or E18.5 (g), a subset in superficial L2 (GFP^+^), projected to iM2. (h) GFP+ IPNs (dots) in S1bfd born at different embryonic times that project to iM2. X axis: age of TM induction; Y axis: depth from pial surface (μm) Asterisk, site of injection (left) and S1_bfd_ area of analysis with high mag insets (right) (b-g). Scale bars = 1mm for brain hemisections, low mag images showing cortical layers = 100μm, high mag images = 20μm. Abbreviations: IS, intersection-subtraction; rAAV, retrograde adeno-associated virus; M2, secondary motor cortex.

### Distinct PN types within the same layer are generated at different times

Our fate mapping in *Tbr2-CreER;Ai14* mice showed that L3 PNs are generated at two non-contiguous embryonic times E14.5 and E16.5 (Figure 2). Since L3 contains both CSPNs and CPNs, we examined whether these two PN subpopulations were differentially generated at the two embryonic times. We performed *rAAV-flp* injections in either ipsilateral DL-Str or contralateral S1 (cS1) in P30 *Tbr2-2A-CreER;IS* mice induced at E14.5 or E16.5, and examined cell labeling patterns in ipsilateral S1_bfd_ (iS1) (Figure 6a,b). Whereas cS1 injection in E14.5-induced mice labeled mostly L3 and L5a (and a few L6) PNs in iS1 (Figure 6c,g; also Figure 4j,n), DL-Str injection in these mice labeled L5a but not L3 PNs, suggesting that L3 PNs born at E14.5 are mostly CPNs but not CSPNs (Figure 6d,g; also Figure 4c,f). On the other hand, cS1 and DL-Str injections in E16.5-induced mice both labeled L3 PNs in iS1 (Figure 6e,f,g; also Figure 4d,f,k,n). Therefore, although E14.5 and E16.5 IP-derived PNs both reside in L3, they represent two different projection types, the former projects to contralateral cortex while the latter projects to both contralateral cortex and the striatum.

**Figure 6:**
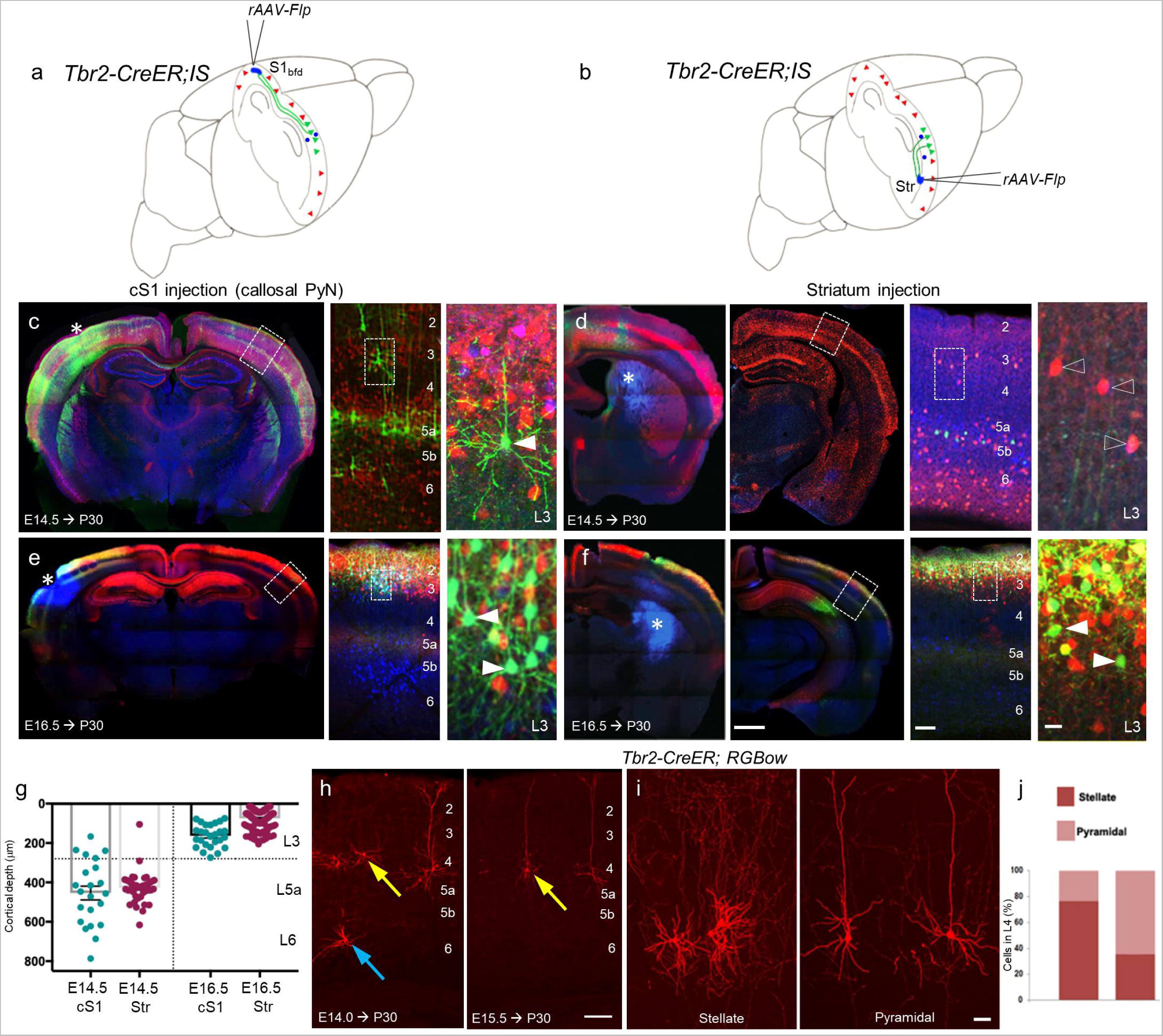
Distinct PN types are generated within Layer 3 and Layer 4 at non-contiguous stages. (a,b) rAAV injection strategy for for fate mapping L3 PNs in S1 projecting to cS1 (a) or DL-striatum (b). (c,d) rAAV-Flp injection (asterisk) in cS1 (c) but not in DL-striatum (d) labeled a subset of GFP^+^ L3 PNs in S1 (open arrowheads, right panels in d) in Tbr2-CreER;IS mice TM-induced at E14.5. (e,f) rAAV-Flp injection (asterisk) in cS1 (e) as well as DL-striatum (f) labeled a subset of GFP^+^ L3 PNs in S1 (arrowheads, right panels in e,f) in Tbr2-CreER;IS mice TM-induced at E16.5. High mag insets for S1_bfd_ in the right panels of c-f. (g) GFP^+^ PNs in S1_bfd_ from injections in cS1 (blue dots) and striatum (red dots) in animals TM induced at E14.5 or E16.5. Graph above the dotted line represents L3. (h) TM induction in Tbr2-CreER;RGBow (also see Fig S3a) at E14.0 (left) or E15.5 (right) sparsely labeled PNs in a pattern similar to the Ai14 reporter (Figure 2). Blue arrows, L6; Yellow arrows, L4 (i) Two distinct morphological PN subtypes, stellate (left) and pyramidal (right), were labeled in L4 of S1_bfd_ cortex in mice TM-induced at E14.0 or E15.5. (j) The majority of E14.0-born L4 PNs are spiny stellate (76.2%, mean + SEM), whereas E15.5-generated L4 PNs are predominantly pyramidal (63.8%, mean + SEM). (n = 6, 3 litters each). Scale bars = 1mm for brain hemisections, low mag images showing cortical layers = 100μm, high mag images = 20μm.

Similar, L4 neurons are also generated from two non-contiguous embryonic times at E14.0 and E15.5 (Figure 2). L4 contains multiple glutamatergic neuron types that mostly project locally^5,56^. To examine whether E14.0 and E15.5 born L4 neurons are the same or different cell types, we examined their morphology using sparse labeling with the *RGBow* reporter (Figure 6h,i; Figure S3). A low dose TM induction in *Tbr2-CreER;RGBow* mice at E14.0 and E15.5 labeled two morphologically distinct populations within L4. Whereas E14.0 induction predominantly labeled spiny stellate cells, characterized by their multipolar dendrites and exuberant vertically projecting axons, E15.5 induction largely labeled pyramidal neurons characterized by prominent apical and basal dendrites and few local axon branches (Figure 6h,i). Quantification of these two morphological types revealed that ~75% of E14.0 born L4 cells were spiny stellate cells, while ~67% of E15.5 born L4 cells were pyramidal neurons (Figure 6j). To further characterize the axon projection of these two L4 subpopulations, we used *rAAV-Flp* injection in *Tbr2-CreER;IS* mice. *rAAV-Flp* injection in cS1 or iM2 of E14.0- and E15.5-induced mice only labeled local L4 neurons but not iS1 neurons, indicating that these L4 neurons are local projecting but not CPN or IPNs (Figure S5). Together, our results demonstrate that IP-derived PNs residing in the same layer that are generated at non-contiguous times represent different PN types. Interestingly, in both L4 and L3, the earlier born PN type occupied a deeper position within the layer compared to the later born PN type.

In summary, by integrating PN birthdating and anatomical analysis of their projection targets and/or morphology, we reveal that that a unifying theme of iNG is the sequential specification of projection-defined PN types, irrespective of their laminar location, by fate-restricted IP temporal cohorts.

## DISCUSSION

Although the cerebral cortex is characterized at the histological level as consisting of dozens of functional areas and multiple cell layers, its fundamental units are the diverse neuronal cell types that assemble cortical circuits^5,6^. Among cortical neuron types, glutamatergic projection neurons (PNs) make up the vast majority, are distributed across layers, and constitute the key elements that construct local circuitry, intracortical processing streams, and cortical output channels^5,6^. Therefore, a satisfying understanding of cortical development needs to explain the origin of diverse PN types, their laminar organization, and ultimately their assembly into cortical circuits^6,10^. While it is now well established that all PNs are generated from radial glial progenitors (RGs) through either direct neurogenesis (dNG) or indirect neurogenesis (iNG) via intermediate progenitors (IPs)^11,17^, the specification and laminal deployment of diverse PN types remain not well understood.

Classic cell birth dating experiments in mice revealed an overall trend of deep-to-superficial level cortical cell migration^21^. The literal interpretation of these results - that “Cells forming successively later in embryonic life migrate outward past their predecessors and come to lie at more and more superficial levels”^21^, has led to the foundational “inside-out model” of corticogenesis^10,23,24^. However, in retrospect, these pioneering studies are substantially limited by the coarse temporal resolution in cell birth dating and the absence of cellular resolution in recognizing different neural progenitor types and diverse neuronal progeny types. Indeed, embryonic tritiated thymidine administration labeled neurons across broad cortical layers^21^, and subsequent higher temporal resolution studies revealed simultaneously-born cells occupying more than one layer as well as substantial overlapping distributions of cells arising with successive cell cycles^25^. Furthermore, the classic studies^21,22^ were carried out long before the discovery of the subpallium origin and tangential migration of GABAergic interneurons^26^; thus the mixed labeling of pallium and subpallium progenitors and their glutamatergic and GABAregic neuronal progeny that migrate along orthogonal routes substantially complicates the interpretation of labeling and deployment pattern of PNs. Third, the findings were made long before the discovery cortical neural progenitors^11,17^, thus did not distinguish the role of IPs and iNG in PN production and laminar deployment. Therefore, to date, direct evidence for a stringent inside-out model of corticogenesis has yet to be experimentally shown, and its biological significance in cortical circuit development remains unclear.

By combining a set of novel genetic tools centered around *Tbr2*, we have achieved fate mapping individual IPs and IP temporal cohorts with the resolution of individual cell division cycles. We further resolve IP-derived and birth-dated PN progeny in terms of their morphology and projection targets - defining features of PN type identity beyond laminar location. We found that individual IPs are fate-restricted to generate a pair of morphologically identical “twin” PNs, most likely of the same anatomical type. Our results suggest that iNG involves the orderly specification of multiple temporal cohorts of fate-restricted IPs, which sequentially as well as concurrently generate numerous subsets of PN progeny, each deployed to a specific layer (Figure 7a). As a result, the laminar deployment of successive IP-derived PN subpopulations substantially deviate from a simple inside-out rule: PNs destined to 2 or 3 non-consecutive layers are generated at the same time, and different PN types of the same layer are generated at non-contiguous times. By integrating cell birth dating with projection mapping, we found that PN birth order largely correlates with their projection pattern, a key determinant of PN type identity (Figure 7b). Our findings therefore suggest that, beyond an overall deep-to-superficial trend of cell migration^21^ and instead of a simple inside-out model of cortical lamination^17,23,24^, the overarching scheme of corticogenesis is the sequential specification and precise laminar deployment of projection-defined PN types, which may contribute to the orderly assembly of cortical output channels and processing streams (Figure 7b). Although our study does not address PNs produced from dNG, iNG generates the large majority of PNs in rodents^24,28,30,31^ and especially in primate^34,38^ and thus represents the dominant mode of corticogenesis.

**Figure 7:**
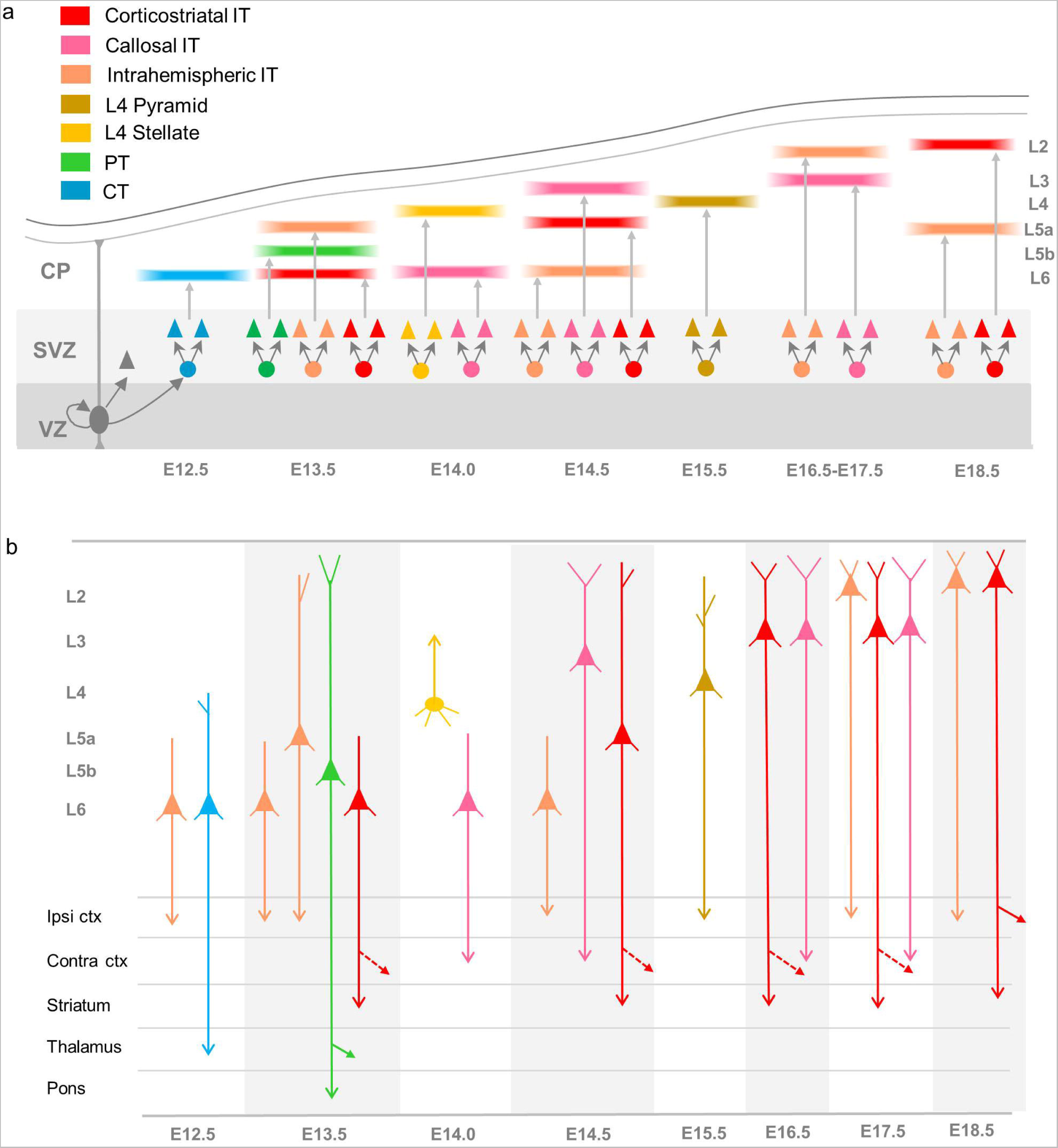
Orderly specification and precise laminar deployment of IP-derived PN types. A model on the role of indirect neurogenesis and intermediate progenitors in corticogenesis. iNG involves the orderly specification of multiple temporal cohorts of fate-restricted IPs, which sequentially as well as concurrently generate numerous subsets of PN progeny, each deployed to a specific layer (a). The laminar deployment of successive IP-derived PN subpopulations substantially deviate from a simple inside-out rule. The overarching scheme of iNG is the sequential specification and precise laminar deployment of projection-defined PN types (b), which may contribute to the orderly assembly of cortical output channels and processing streams.

As all IPs are generated from RGs, the sequential specification of temporal cohorts of fate-restricted IPs most likely results from the lineage progression of RGs. An open question is the relationship between dNG- and iNG-derived PNs born at the same time, addressing which would require a method to simultaneously birthdate RG-and IP-derived PNs. The fate-restriction of individual cortical IPs to give rise a pair of twin PNs should not be taken for granted, as in *Drosophila,* each iNG division mostly leads to the generation of two distinct cell types^57,58^. Therefore, whereas the specification of multiple fate-restricted IP temporal cohorts diversifies PN production, the amplification of IP numbers within each cohort and the production of two identical PNs from each IPs expand the production of each PN type. Notably, a recent study demonstrates that *Tbr2^+^* IPs at different times of corticogenesis co-express neuronal identity genes characteristic to postmitotic PNs born at the corresponding time^59^, supporting the notion that IPs are transcriptionally poised to generate specific PN types. Our genetic fate mapping tools will facilitate targeted transcriptomic and epigenomic analysis of the developmental trajectory of iNG, from fate-restricted IPs to their progeny PN types, to discover the underlying genetic regulatory programs.

A previous study using a *Tbr2-CreER* driver line in which a *CreER* cassette was inserted at the translation initiation codon, inactivating one *Tbr2* allele^60^, provided evidence for IP laminar fates that correspond to upper cortical layers during early cortical neurogenesis^32^. However, the analysis of fate mapped PN progeny was carried out at P0.5, when distinct cortical layers are not yet fully formed^61,62^ and the molecular markers assayed do not yet differentiate major PN types^53,63,64^; thus the study only broadly parsed PNs into upper and lower layer populations and did not resolve the relationship between IP temporal cohorts and PN projection types. A follow up study carried out clonal analysis with *MADM* at E11.5 and E12.5, but the analysis of PN progeny was again performed very early at E18.5, before their laminar fate could be fully ascertained^65^. In both studies, the inactivation of one *Tbr2* allele might impact IP differentiation, as *Tbr2* mutations have been implicated neurodevelopmental disorders^66,67^. As such, these previous results are not directly comparable to our findings.

The precise laminar deployment of concurrently generated PNs to non-consecutive layers and non-contiguously generated PNs to sublaminae of the same layer suggests highly stringent molecular mechanisms instilled in the specified PN types that guide their migration and settlement. While our retrograde labeling experiments begin to reveal the broad projection class identities of IP-derived PNs (e.g. CT, PT, CSPN, CPN), single cell RNA sequencing^9^ and spatial transcriptomic analysis^68^ of fate mapped samples will resolve these PN types at a higher granularity. Such studies promise to uncover the molecular mechanisms that underlie not only their precise laminar deployment but also their specific projection patterns and possibly synaptic connectivity, which together contribute to the assembly cortical circuits and brain networks.

## Acknowledgments

We thank Debra L. Silver and Dmitry Velmeshev for comments on the manuscript. We thank Hayley Porter, Chris Xu, Ricardo Raudales and Ankit Sood for help with quantification related to Figures and the CSHL Microscopy shared resource. This work is supported by NIH grant U19MH114823-01 (Z.J.H.). ZJH is supported by a NIH Director’s Pioneer Award 1DP1MH129954-01. DH was supported by a Human Frontier Science Program long-term fellowship LT000075/2014-L and a NARSAD Young Investigator grant no. 26327. J.M.L. was supported by a NRSA F30 Medical Scientist Predoctoral Fellowship 5F30MH108333. B.-S.W was supported by a NRSA Postdoctoral Fellowship NIH5F32NS096877-03.

## Author contributions

Conceptualization: Z.J.H., D.H. and J.M.L; experimental design: Z.J.H., D.H., and J.M.L.; experimental investigation: D.H., J.M.L., W.G. (fate mapping, immunohistochemistry, imaging, and quantification), J.M.L., D.H., B.-S.W., and W.G. (stereotaxic injections); writing, reviewing, and editing: Z.J.H., D.H.

## METHODS

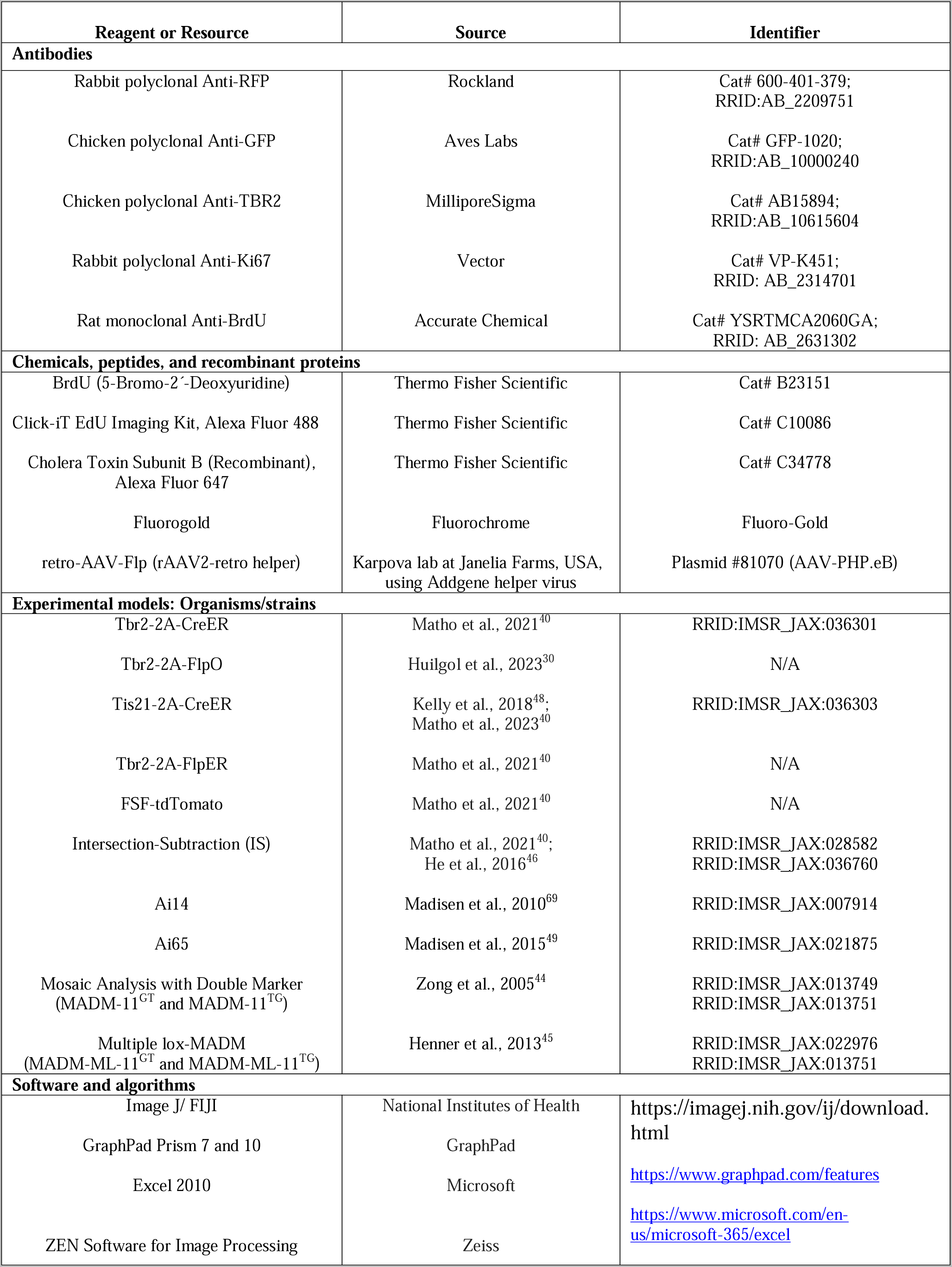
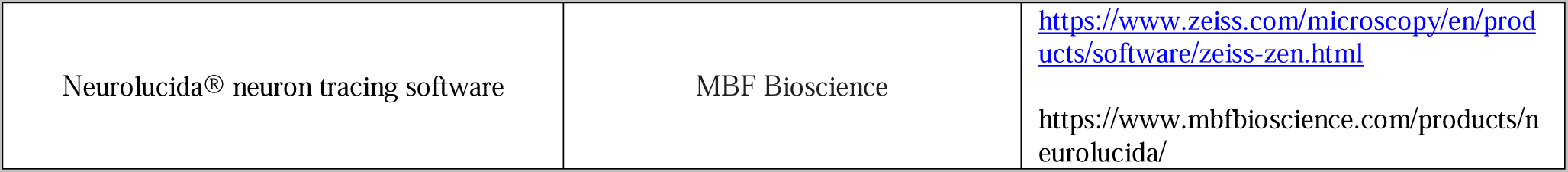

### Experimental model and study participant details

We have used embryonic and adult genetically targeted mice (Mus musculus) for this study. In all experiments using P30 animals, male and female mice are analyzed in equal or near-equal numbers. We did not observe any sex-related differences in our results using P30 animals. No sex determination was performed for embryonic analysis. All genetic knock-in mice have been backcrossed 6 generations to a Swiss-Webster background.

All adult histology experiments were performed between postnatal day (P)30-P45. Retrograde injections in Figures 3-6 were analyzed at ~P45. We harvested embryos from embryonic day (E)10.5-E16.5 for Tbr2-2A-CreER mice. Embryonic and postnatal ages are denoted in Figures, Figure legends and relevant places in the Results.

We have used the following knock-in mice in this study: Driver lines include Tbr2-2A-CreER, Tis21-2A-CreER, Tbr2-2A-FlpER, Tbr2-2A-FlpO. Reporter lines include Ai14, MADM, Ai65 and Intersection-Subtraction (IS) (See Key Resources Table). All fate-mapping and birth-dating experiments were conducted by combining one or more of the driver mouse lines with one of the reporter lines and/or Swiss-Webster mice (Taconic).

Mouse related experimental procedures were approved by the Institutional Animal Care and Use Committee (IACUC) of Cold Spring Harbor Laboratory and Duke University in accordance with NIH guidelines.

### Tamoxifen induction and PN birth dating

Tamoxifen (TM) (T5648, Sigma) was prepared by dissolving the powder in corn oil (20 mg/ml) and either applying a sonication pulse for 60s or constant magnetic stirring overnight at 37 °C. A 50 mg/kg dose was administered by oral gavage at the appropriate embryonic age for Tbr2-CreER; Ai14, Tbr2-CreER; IS, Tbr2-CreER; MADM, Tbr2-CreER; RGBow and Tis21-CreER;Tbr2-FlpER;Ai65 animals. For MADM experiments, 300mg/kg TM induction dose and for RGBow experiments, 2mg/kg dose was orally administered.

5-bromo-2’-deoxyuridine (BrdU) Thermo Fisher Cat #B23151) was injected intraperitoneally (100mg/kg) simultaneously with TM induction in pregnant dams for birth dating.

EdU (100mg/kg) (Thermo Fisher, Cat no. C10086) was injected intraperitoneally 24 hours following TM induction to determine any non-IP persisting progenitor types.

### Immunohistochemistry

Adult mice (>P28) were anaesthetized (using Avertin or isoflurane) and transcardially perfused with saline followed by 4% paraformaldehyde (PFA) in 0.1 M phosphate buffer. Brains were rinsed three times in PBS after post-fixation in PFA and sectioned at a 65-70μm thickness with a Leica VT1000S vibratome. Embryo heads were collected in PBS and fixed in 4% PFA for 4-5h at room temperature, rinsed three times with PBS, equilibrated in 30% sucrose-PBS, frozen in OCT compound (Thermo Fisher, Cat no. 23-730-571) and cut on a cryostat (Leica, CM3050S) at 25-35μm coronal sections.

Sections were treated with a blocking solution (10% normal goat serum and 0.2% Triton-X100 in 1X PBS) for 2h, then incubated overnight at 4°C with primary antibodies diluted in the blocking solution. Sections were washed three times in PBS and incubated for 2h at room temperature with corresponding secondary antibodies, Goat or Donkey Alexa Fluor 488, 594 or 647 (1:500, Life Technologies) and DAPI to label nuclei (1:1000 in PBS, Life Technologies, 33342), NeuroTrace 435/455 (1:300, Thermo Fisher) or NeuroTrace 640/660 (1:300, Thermo Fisher) diluted in PBS with 0.2% triton X-100 for 2-4 hours at room temperature. Sections were washed three times with PBS and dry-mounted on slides using Fluoromount-G (SouthernBiotech, 0100-01) mounting medium.

For BrdU immunostaining, before the blocking step, sections were incubated with 2N HCl at 37C for 45 min followed by neurtralization in 0.1M sodium tetraborate (pH 8.5) for 10 minutes at roo temperature. Sections were again rinsed in PBS and subsequently mounted in Fluoromount-G (SouthernBiotech).

#### Primary Antibodies

Anti-RFP (1:1000, Rockland Pharmaceuticals, 600-401-379), anti-GFP (1:1000, Aves, GFP-1020), anti-BrdU (1: 300, Accurate Chemical, YSRTMCA2060GA), anti-TBR2 (1:250, MilliporeSigma AB15894), anti-Ki67 (1:300, Vector, Cat# VP-K451).

Click-iT EdU Imaging Kit, Alexa Fluor 488 (Thermo Fisher, Cat no. C10086) was used for processing adult brains from mice that were administered with EdU.

### Stereotaxic Injections

Adult mice, P28 or older were injected under 2% isoflurane in oxygen anesthesia with 0.41/min airflow on a stereotactic frame (Kopf instruments, 940 series). Craniotomy was performed with a dental drill. Preemptive analgesics, 5mg/kg ketoprofen and 0.5mg/kg dexamethasone, were administered subcutaneously before the surgery. Lidocaine (2–4 mg/kg) was applied intra-incisionally. Injection coordinates were selected based on Allen Brain Atlas measurements for S1bfd cortex and projection targets from that region. Tracers were loaded into glass capillaries with 1.17mm inner diameter, 1.50mm outer diameter and injected using a picospritzer (Parker) with a duration of 30ms at a frequency of 1Hz. Tracers used include retro-AAV-Flp (Karpova Lab at Janelia Farms, USA) and FluoroGold (FG) (FluoroChrome). Viruses were incubated for 2 weeks prior to sacrificing the animal. A pulled glass pipette tip of 20–30 μm containing CTB647 (ThermoFischer Scientific, C34778) or retrograde AAV (Addgene, AAV-PHP.eB) and FG was lowered into the brain. A 500nl (CTB or FG) and/or 300-400nl (retroAAV) volume was delivered at a 30nl/min using a Picospritzer (General Valve Corp); to prevent backflow, the pipette was maintained in place for 10 min prior to retraction. The incision was sutured with Tissueglue (3M Vetbond), following which mice were kept warm at 37°C until complete recovery.

### Imaging

All imaging was done using Zeiss LSM 710, 780 or 900 (CSHL St. Giles Advanced Microscopy Center, Duke University Light Microscopy Core Facility and our laboratory) fluorescence confocal microscopes using objectives, 5x for tilescan, 10x or 20x for z-stacks. For embryos, high magnification images were obtained using 63x oil objective. To determine colocalization in adult mouse brains, confocal Z-stacks were obtained centered in S1bfd, using a 20x objective. We manually determined colocalization for the desired markers by looking in individual Z-planes using ImageJ/FIJI software. For embryonic analysis (Figures 1, S1), high-magnification insets are not always maximum intensity projections. To observe the morphology of IPs and quantification of colocalization, only a few sections from the Z-plane in low-magnification images have been projected in the high-magnification images.

### Quantification

All quantifications were performed by two individuals (one blinded). Statistics and plotting of graphs were done using GraphPad Prism 7, 10 and Microsoft Excel 2010.

For embryonic quantifications, we counted 70-200 cells depending on the extent of labeling from at least 5 embryonic brains across 2 litters. For colocalization experiments with TBR2 and Ki67 in embryonic brains, DAPI (not shown) was used to identify cells.

For P30 brains, all quantifications were done at primary somatosensory area, barrel field (S1bfd) (Bregma 1.0-2.0) according to the Mouse Brain in Stereotaxic Coordinates 4th Edition atlas (Franklin and Paxinos). Experiments using Tbr2-CreER; Ai14 with BrdU and TM co-injections (Figure 2), laminar positions of cells was detected using an ImageJ/FIJI plug-in for automated cell depth measurement by extrapolating X-Y coordinates of each cell from the pial surface. NeuroTrace™ 640/660 fluorescent Nissl (Cat no. N21483) was also used to determine cell packing density and the layer positions (not shown).

Counting of *Tbr2-CreER;MADM* brains was completed using Neurolucida software.

#### Quantification of rAAV-Flp injection experiments

Automated counting of cells labeled by rAAV-Flp-containing virus injected in Tbr2-CreER;IS mice was completed using ImageJ/FIJI from confocal images. Automated cell depth measurements were made by extrapolating X and Y coordinates of each cell and measuring the shortest distance between that point and the contour of the section, representing distance from the pial surface.

