## Supplemental Information for "Orderly specification and precise laminar deployment of cortical glutamatergic projection neuron types through intermediate progenitors"

**SUPPLEMENTARY FILE**

**Contains Figures S1-S5 and their figure legends, Supplementary video S1 legend.**

Figure S1_Huilgol

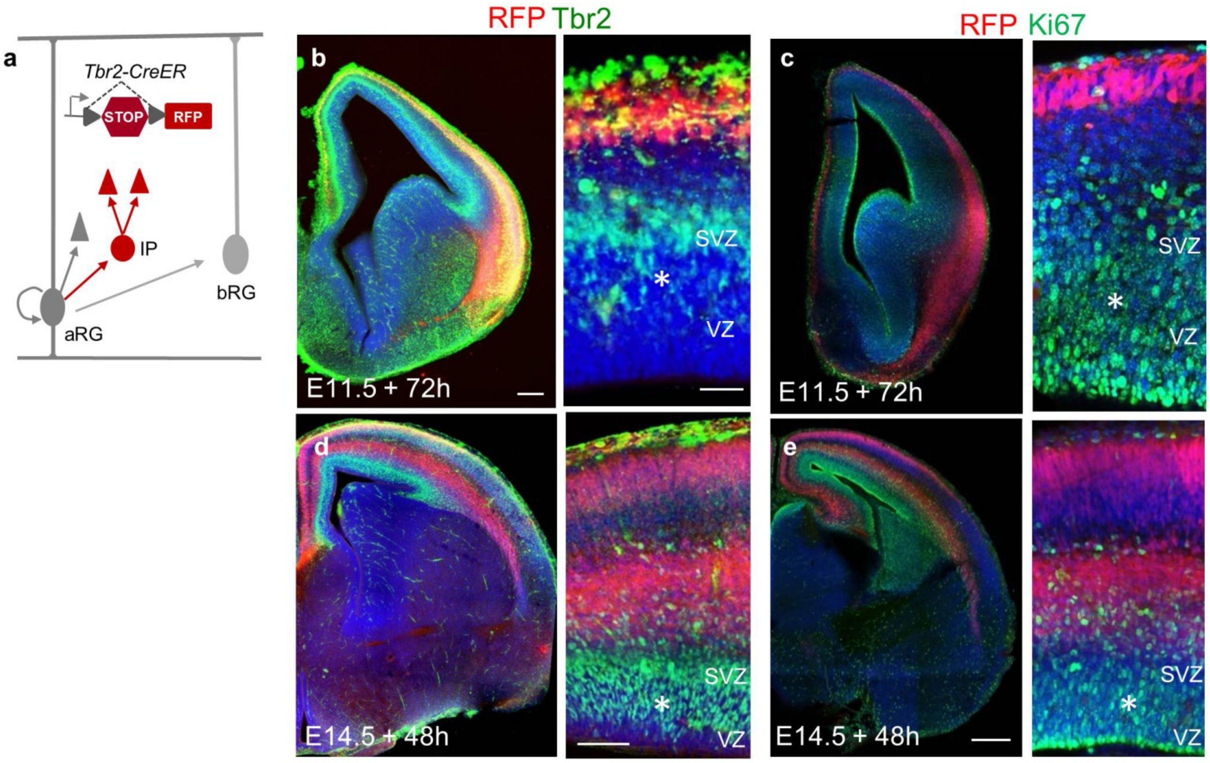


**Figure S1: *Tbr2-creER* does not label basal radial glia (Related to Figure 1)**

(a) Schematic showing that *Tbr2-CreER;Ai14* labels IP (red) but not basal radial glia (grey).

(b,c) 72 hours after TM induction at E11.5 shows no colocalization of $RFP^{+}$ cells in VZ/SVZ (asterisk) with TBR2 (b) and the cell cycle marker Ki67 (c).

(d,e) 48 hours after TM induction at E14.5 also shows no overlap (asterisk) between RFP and TBR2 (d) or Ki67 (e). Magnified images (right). (n=4, 2 litters). Scale bars = 100□m. Abbreviations: VZ, ventricular zone; SVZ, subventricular zone; aRG, apical radial glia; bRG, basal radial glia; IP, intermediate progenitor.

Figure S2_Huilgol

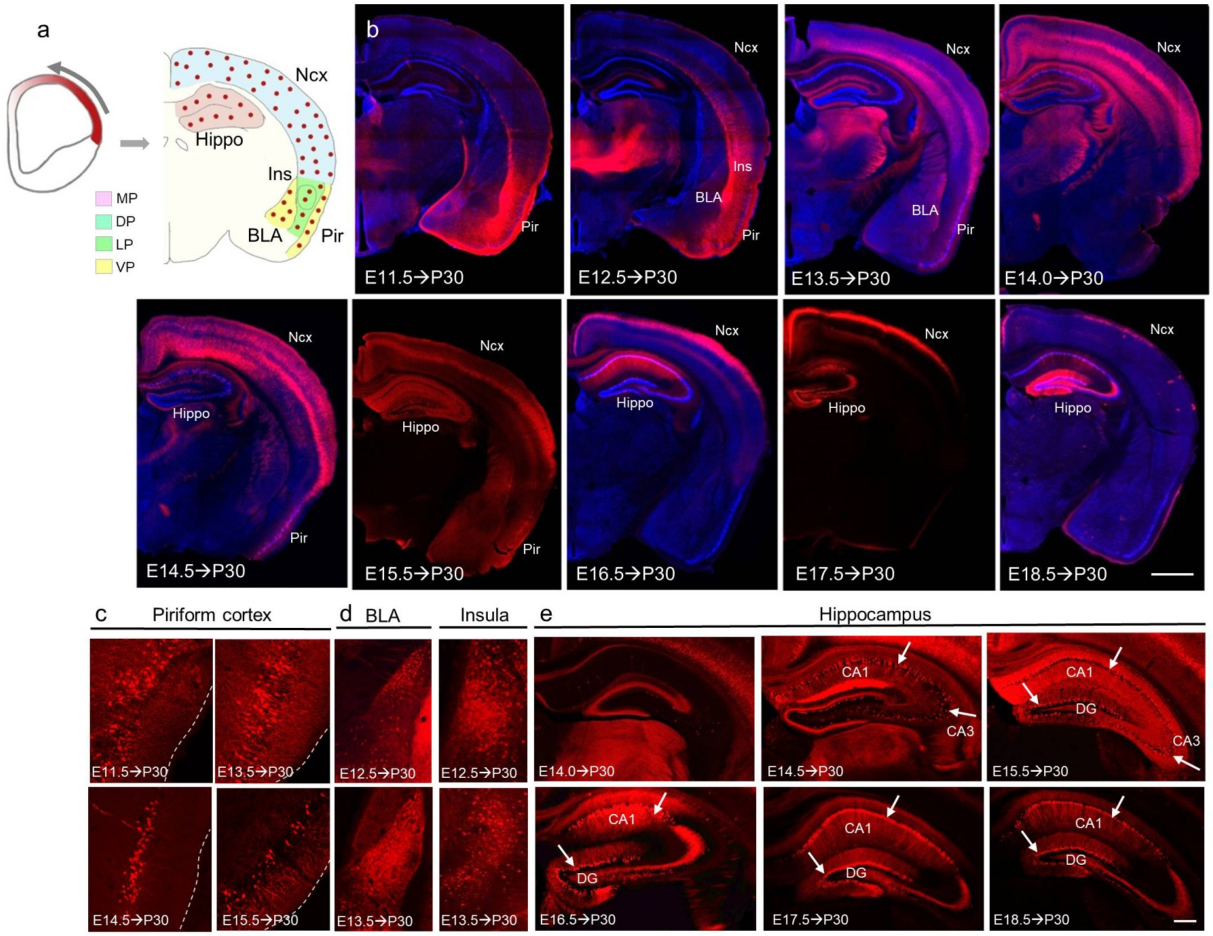


**Figure S2 (Related to Figures 1 and 2): Tbr2$^+$ IPs generate PNs in cortical structures in a lateral-to-medial temporal gradient**

(a) Schematic summarizing Tbr2$^+$ IPs (red) labeled in a lateral-high to medial-low gradient in the embryonic pallial neuroepithelium (left) which generate PNs (red dots) in different cortical structures derived from the four pallial subdivisions (right); MP (Medial pallium; pink), DP (Dorsal pallium; blue), LP (Lateral pallium; green); VP (Ventral pallium; yellow).

(b) The lateral-to-medial generation of PNs from VP and LP-derived structures such as BLA, Pir beginning at E11.5-E12.5, followed by DP-derived neocortex that begins at E11.5 and through E18.5. iNG in MP-derived hippocampus occurs predominantly from mid-to-late stages (E14.5 to

E18.5) of corticogenesis. This lateral-to-medial temporal gradient of iNG is reminiscent of *Tbr2* gradient in the embryonic SVZ.

(c) Temporal pattern of iNG in piriform cortex beginning at E11.5 and ceasing by ~E15.5.

(d) iNG in BLA (left) and insular cortex (right) at E12.5 and E13.5

(e) iNG in the hippocampus begins at ~E14.5 (when CA1 and CA3 PNs are born) with sparsely produced DG cells, followed by the generation of PNs for all three subfields at E15.5. From E16.5-E18.5, iNG generates PNs for the CA1 as well as the DG. Scale bars = 1mm for brain hemisections, high mag images = 100□m. Abbreviations: Ncx, neocortex; Hippo, hippocampus, Ins, insular cortex; Pir, piriform cortex; BLA, basolateral amygdala; MP, medial pallium; DP, dorsal pallium; LP, lateral pallium; VP, ventral pallium; CA, cornu ammonis; DG, dentate gyrus.

Figure S3_Huilgol

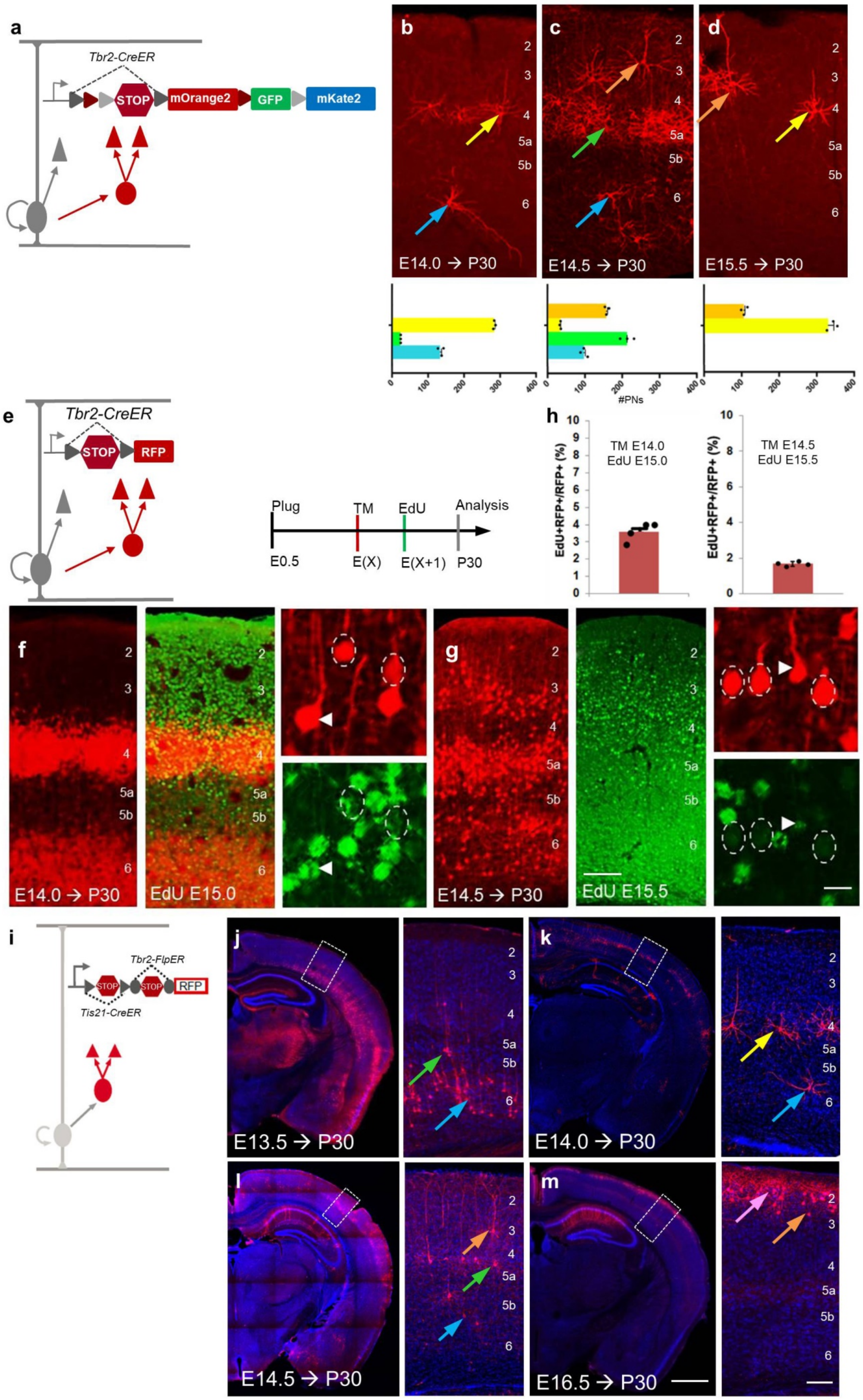

**Figure S3: PNs in multiple non-consecutive layers are generated at the same time from IPs (Related to Figure 2)**

(a) Schematic using *Tbr2-CreER* and ther *Cre*-dependent RGBow reporter (also see Figure 6h,i) to sparsely label IPs and their PN progeny.

(b-d) High-magnification images of P30 *Tbr2-CreER;RGBow* coronal hemisections (top) TM-induced at E14.0 (b), E14.5 (c) and E15.5 (d), and quantified (bottom). (n=3, 2 litters).

(e) Genetic strategy to test the presence of transit-amplifying IPs using *Tbr2-CreER;Ai14* (left) and EdU administration 24 hours after TM induction (right).

(f-h) P30 brains TM-induced at E14.0 (f) and E14.5 (g) showed low presence of transit amplifying IPs. High magnification images (right) show colocalized (arrowheads) and non-colocalized (dashed circles) cells, quantified (h) (n=4, 2 litters).

(i) Genetic birth-dating scheme using *Tis21-CreER;Tbr2-FlpER* bred with an intersectional *Cre-* and *Flp*-dependent reporter, *Ai65*.

(j-m) TM induction at E13.5 (j), E14.0 (k), E14.5 (l) and E16.5 (m). (n=3, 2 litters at each age). L6 - blue; L5 - green; L4 - yellow; L3 - orange (arrows and bar graphs). Scale bars = 1mm for brain hemisections, low mag images showing cortical layers = 100□m, high mag images = 20□m. Abbreviations: TM, tamoxifen; EdU, 5-Ethynyl-2′-deoxyuridine.

Figure S4_Huilgol

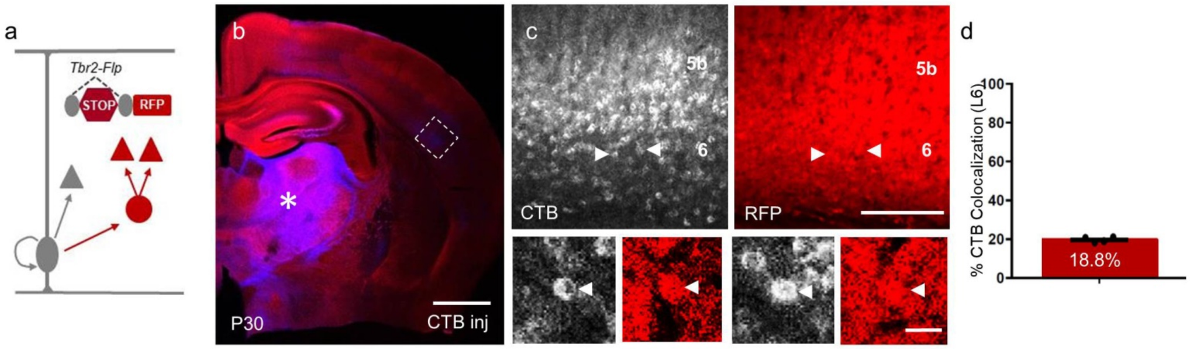


**Figure S4: iNG produces smaller proportion of L6 CTs (Related to Figure 3)**

(a) Schematic showing *Tbr2-FlpO* bred with a *Flp*-dependent *Rosa26-frt-STOP-frt-RFP* reporter, which labels all iNG-derived PNs.

(b) Coronal hemisection showing a broad thalamus injection of $CTB^{647}$ in P30 *Tbr2-FlpO;FSF-RFP* mice (asterisk).

(c) (top) CTB colocalization with $RFP^+$ cells in L6 (arrowheads). (bottom) High magnification showing two representative examples of $RFP^+$ cells colocalized with CTB (arrowheads).

(d) CTB and $RFP^+$ PNs showed 18.8% colocalization in L6 throughout all cortical areas.

Scale bars = 1mm (b), low mag images showing cortical layers = 100□m, high mag images = 20□m (c). Abbreviations: CTB, cholera toxin subunit B.

Figure S5_Huilgol

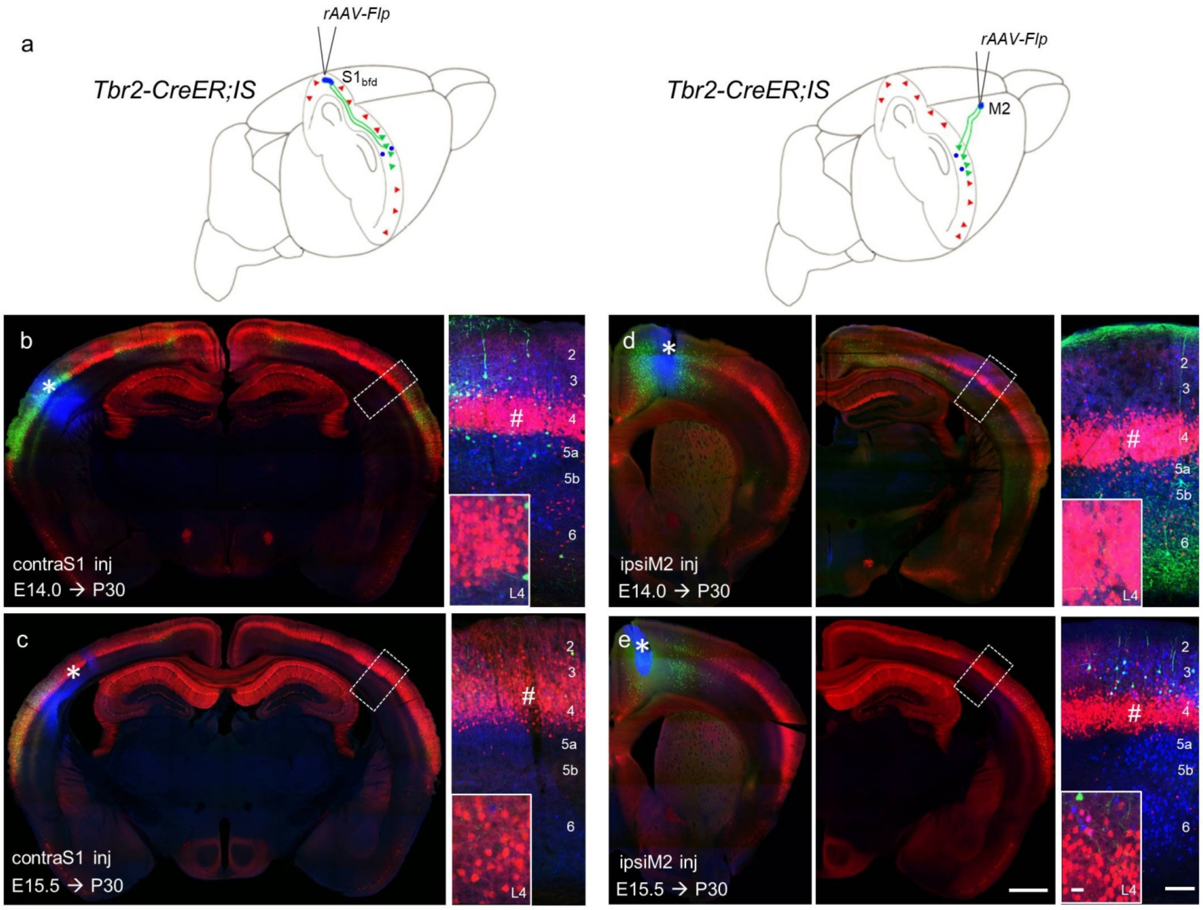


**Figure S5: iNG generates locally projecting Layer 4 PNs (Related to Figure 6)**

(a) Same strategy as in Figure 3a to examine locally projecting L4 neurons *Tbr2-CreER;IS* mice.

(b,c) *rAAV-Flp* injection into the cS1 (asterisk, left) in mice TM-induced at E14.0 (b) or E15.5 (c). Analysis in $S1_{bfd}$ shows no $GFP^+$ PNs in L4 (hashtag and high mag insets, right panels).

(d,e) M2 injection (asterisk, left) in mice TM-induced at E14.0 (d) or E15.5 (e) did not label $GFP^+$ PNs in L4 in ipsilateral S1 (hashtag and high mag insets, right panels).

High magnification insets show L4 (b-e). Scale bars = 1mm for brain hemisections, low mag images showing cortical layers = 100□m, high mag images = 20□m.

**Supplementary video 1: Twin PNs generated simultaneously in different cortical structures**

3D reconstruction of P30 brain sections (70μm thickness) from *Tbr2-CreER;MADM*, TM-induced at E14.0 using Neurolucida (MBF Biosciences). Red-Green (RG) PN pairs observed in cortical structures including the neocortex, piriform cortex, basolateral amygdala (purple) and the hippocampus (green). Striatum (blue). Each dot represents a PN, red dot = RFP$^+$ cell, green dot = GFP$^+$ cell. A total of 18 RG pairs observed of total 24 red and 24 green cells.
